# Dendritome Mapping Unveils Spatial Organization of Striatal D1/D2-Neuron Morphology

**DOI:** 10.1101/2024.10.24.619934

**Authors:** Chang Sin Park, Ming Yan, Muye Zhu, Masood A. Akram, Nicholas N. Foster, Andrew Bennecke, Christopher Choi, Karl Marrett, Keivan Moradi, Jason Y. Zhang, Gabrielle Magat, Sumit Nanda, Raymond Vaca, Kathleen Wijaya, Jedrick Regala Zablan, Sebastian Lee, Cassidy Song, Mary Jasmine Lara, Madeline Louie, Jason Cong, Yongsoo Kim, Giorgio A. Ascoli, Peter Langfelder, Daniel Tward, Hong-Wei Dong, X. William Yang

**Affiliations:** Center for Neurobehavioral Genetics, Jane and Terry Semel Institute of Neuroscience and Human Behavior, University of California, Los Angeles, Los Angeles, CA, USA; Dept. Psychiatry and Biobehavioral Science, David Geffen School of Medicine, University of California, Los Angeles, Los Angeles, CA, USA; UCLA Brain Research and Artificial Intelligence Nexus, Department of Neurobiology, David Geffen School of Medicine, University of California, Los Angeles, Los Angeles, CA, USA; Departments of Computational Medicine and Neurology, University of California, Los Angeles, Los Angeles, CA, USA; Department of Computer Science, University of California, Los Angeles, Los Angeles, CA, USA; Department of Neural and Behavioral Sciences, The Pennsylvania State University, Hershey, PA, USA; Center for Neural Informatics, Structures and Plasticity, Bioengineering Department and Krasnow Institute for Advanced Study, George Mason University, Fairfax, VA, USA

**Author notes:** Co-corresponding authors. Emails: X.W.Y.; H.-W. D.; D.T.

## Abstract

Morphology is a cardinal feature of a neuron and mediates its functions, but profiling neuronal morphologies at scale remains a formidable challenge. Here we describe a generalizable pipeline for large-scale brainwide profiling of dendritic morphology of genetically-defined single neurons in the mouse brain. We generated a dataset of 3,762 3D-reconstructed and reference-atlas mapped striatal D1- and D2-medium spiny neurons (MSNs). Integrative morphometric analyses reveal distinct impacts of D1/D2 and anatomical locations on MSN morphology. To analyze striatal regional features of MSN dendrites without prior anatomical constraints, we assigned MSNs to a lattice of cubic boxes in the reference brain atlas, and summarized morphometric representation (“eigen-morph”) for each box and clustered boxes with shared morphometry. This analysis reveals 6 modules with characteristic dendritic features and spanning contiguous striatal territories, each receiving distinct corticostriatal inputs. Finally, we found aging confers robust and concordant dendritic length and branching defects in D1/D2-MSNs, while Huntington’s disease (HD) mice exhibit MSN-subtype and striatal regional specific pathology. Together, our study demonstrates a systems-biology approach to profile dendritic morphology of genetically-defined single-neurons; and defines novel striatal D1/D2-MSN morphological territories and aging- or HD-associated pathologies.

## INTRODUCTION

Ramón y Cajal was the first to recognize that a fundamental understanding of the brain requires in-depth knowledge of its basic building block, individual neurons^1^. The U.S. BRAIN Initiative Cell Census Network (BICCN)^2,3^ and other large-scale efforts^4^ endeavored to collect and integrate salient multimodal properties of single neurons (e.g., transcriptome/epigenome, morphology, electric properties, and connectivity) in order to build a taxonomy of the neuronal cell types for the mammalian brain. Currently, single-cell transcriptomes and epigenomes have the highest scalability and enable the identification of >5000 molecularly distinct neuronal clusters^3,5^. However, the morphological diversity of mammalian neuronal cell types, including that of the molecularly- or genetically-defined neurons, remains poorly understood.

Traditional methods to assess neuronal morphology involve sparse labeling of neurons, such as histochemical (e.g., Golgi) staining, manual dye microinjections, or genetic sparse labeling, followed by a laborious process from imaging to manual reconstruction^6,7,8^. Due to the high technical, equipment, labor demands, and low throughput, morphological analyses of single neurons are infrequently employed or used only at a limited scale, e.g. up to dozens of neurons per study^9^. To date, only a handful of studies labeled, imaged, and reconstructed the morphologies (dendrites and/or axons) of over 1000 single neurons in mammalian brains^10,11,12,13,14,15,16^. These studies represent significant advances in scales compared to traditional methods of neuronal morphological studies. However, the approaches in these studies also have limitations in scalability and a lack of broader adoption in the field due to the use of specialized (and not readily available) imaging equipment (e.g., fMOST, STPT), and technically demanding skills (e.g., high-throughput patch clamp recordings). Thus, it remains a critical need to develop streamlined, readily democratizable, and scalable pipelines, to profile neuronal morphology in the mammalian brain.

The mammalian striatum (also called the caudate/putamen) is critical for controlling volitional behavioral output including motor performance, motor skill and habit learning, and higher cognitive function such as decision making and language^17^; its dysfunction or degeneration is implicated in brain disorders ranging from movement disorders^18,19^ to psychiatric disorders and drug addiction^20,21,22^. In the striatum, about 90% of the neurons are medium spiny neurons (MSNs, also known as spiny projection neurons or SPNs), which are divided into direct (D1) and indirect (D2) pathway types based on the expression of dopamine D1- and D2-receptors^23,24,25^ and distinct efferent targets^18,26^. The MSN dendrites are integrators of diverse axonal inputs from the cortex, thalamus, dopaminergic neurons in the midbrain, and other afferents and local neuronal inputs^27^. MSN dendrites are known to be plastic, and their morphologies can change in response to dopamine signaling or dopamine depletion^28,29,30^, chronic stress^31^, and neurodegenerative disease such as Huntington’s disease (HD)^32,33,34^. Despite the importance of MSN dendritic morphology, to date only about 3700 MSN dendrites have ever been reconstructed and publicly archived, with an average of about 64 reconstructed neurons per study (data from NeuroMorpho.Org version 8.6.54). Moreover, our knowledge of differential D1- and D2-MSN dendritic morphologies are based on a few studies with dozens of reconstructed neurons each^35,36^. Given that each mouse striatum has about 1.7-1.9 million MSNs^37^, it is evident that the organization principles of the D1- and D2-MSN dendrites throughout the striatum remain largely unknown.

In this study, we developed a novel, generalizable and complete pipeline to analyze the dendrites of genetically-defined single neurons. We have labeled, imaged, and digitally reconstructed the nearly complete dendritic morphologies for over 3500 D1- and D2-MSNs. We developed a novel neuronal reconstruction and morphometry analysis pipeline, with the latter incorporating network biology tools used in omics analyses (i.e., WGCNA)^38^. Our study uncovers the organizational principles of MSNs based on their D1/D2 genetic types and striatal anatomical territories. Our study also reveals the impact of aging and the mutant Huntingtin (the causal gene in Huntington’s disease) in altering D1- and D2-MSN morphologies. Together, our study demonstrates the importance of both establishing a reference MSN dendritome and then applying it to study brain biology and diseases.

## RESULTS

### A generalizable pipeline for large-scale morphological studies of genetically-defined single neurons in the mouse brain

To enable the large-scale study of the morphology of genetically-defined single neurons, we developed a novel pipeline and conducted a proof-of-concept study of over 3500 genetically-defined D1- and D2-MSNs at postnatal day 56 (P56) and 12 months of age (Fig. 1a). To stochastically and brightly label genetically-defined single neurons, we employed MORF3 mice^8^, which confer Cre-dependent brainwide genetic sparse labeling of single neurons. MORF3 reporter is a membrane-bound tandem “spaghetti monster” reporter V5 reporter with 20 V5 epitopes (td-smFPV5-F), and hence it enables extremely high signal-to-noise labeling of single-neuron dendrites and axons^8^. To label striatal D1- and D2-MSNs at low sparsity to facilitate full dendritic morphology visualization, we crossed MORF3 with CamK2a-CreER^39^ and D1-TdTomato^40^ followed by low tamoxifen induction, which resulted in labeling of only dozens to hundreds of MSNs per brain (Fig. 1a). We collected MSN morphological data from 5 P56 wildtype (WT) control brains and 7 brains from the 12-month (12m) WT and HD-Q140 dataset.

**Fig. 1.**
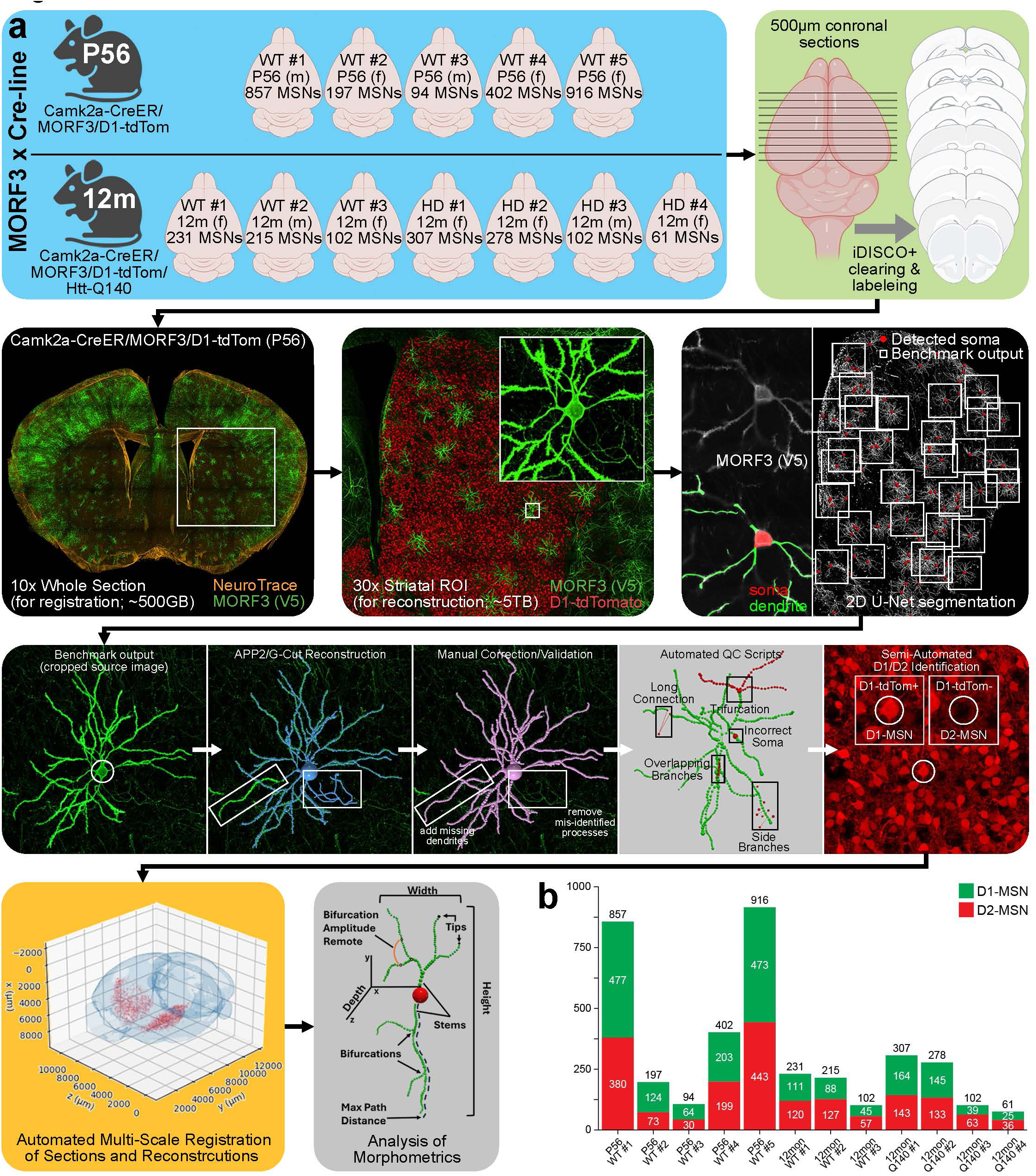
Summary of data generation for P56 and 12mon WT and HD (Q140) striatal MSN dendritome. **a**, Pipeline for high-resolution imaging, semi-automated dendritic reconstruction, automated CCF-registration, and morphometric analysis of MORF3-labeled striatal MSNs in thick-sectioned tissue from P56 and 12mon Camk2a-CreER/MORF3/D1-tdTom mice. **b**, Total number of fully validated MSN reconstructions (with total number of identified D1- and D2-MSNs) from P56 (n=5) and 12mon (WT, n=3; Q140, n=4) brains.

Although MORF3/Cre mice label both dendrites and axons^8,12,41^, we focused this study on the mapping of neuronal dendrites (i.e., “dendritome mapping”) primarily due to the relative scalability of dendritic, but not axonal, digital reconstruction and manual validation. To image relatively complete MSN dendrites, we applied iDISCO+ clearing of 500 µm-thick serial coronal brain sections^8,42,43^, followed by immunolabeling to detect MORF3 (anti-V5) and TdTomato reporters (Fig. 1a), and fluorescent Nissl staining (NeuroTrace) to facilitate brain image registration. The cleared, serial sections were imaged with a DragonFly confocal microscope (Andor)^8^. The former was imaged for the purpose of registration to the reference atlas, i.e., Allen Mouse Brain Common Coordinate Framework or CCFv3^44,45^; and the latter was imaged for dendritic reconstruction of identified D1- and D2-MSNs. We have previously shown that 500 µm section thickness is more likely to capture the complete dendritic morphology of MSNs^8^ compared to the 100-350 µm sections used in the most previous studies^46^.

Our image processing and analysis pipeline includes a series of existing and novel algorithms and computational tools. First, for segmentation of the images, we developed a 2D U-Net classifier that was trained to identify the somas and dendritic processes of individually labeled neurons to facilitate reconstructions. Second, we incorporated in the pipeline G-Cut^47^ to separate any overlapping processes from adjacent neurons and semi-automated reconstruction program’ APP2^48^. Third, we developed a new script called “Benchmark Manager” for isolation of individual neuronal reconstructions and cropping the associated neuronal source image to facilitate manual validation in neuTube^49^. Fourth, we trained a machine learning classifier to automatically identify the presence or absence of D1-tdTomato signal in each labeled D1- or D2 MSNs, respectively (Fig. 1a). Another innovative step is a new multi-scale registration algorithm to map thick brain section images to the CCFv3 (Fig. 2). Lastly, for each validated MSN reconstruction, we extracted 28 morphometric features with L-Measure^50,51^ and 3 with the TREES Toolbox^52^ (Fig. 1a).

**Fig. 2.**
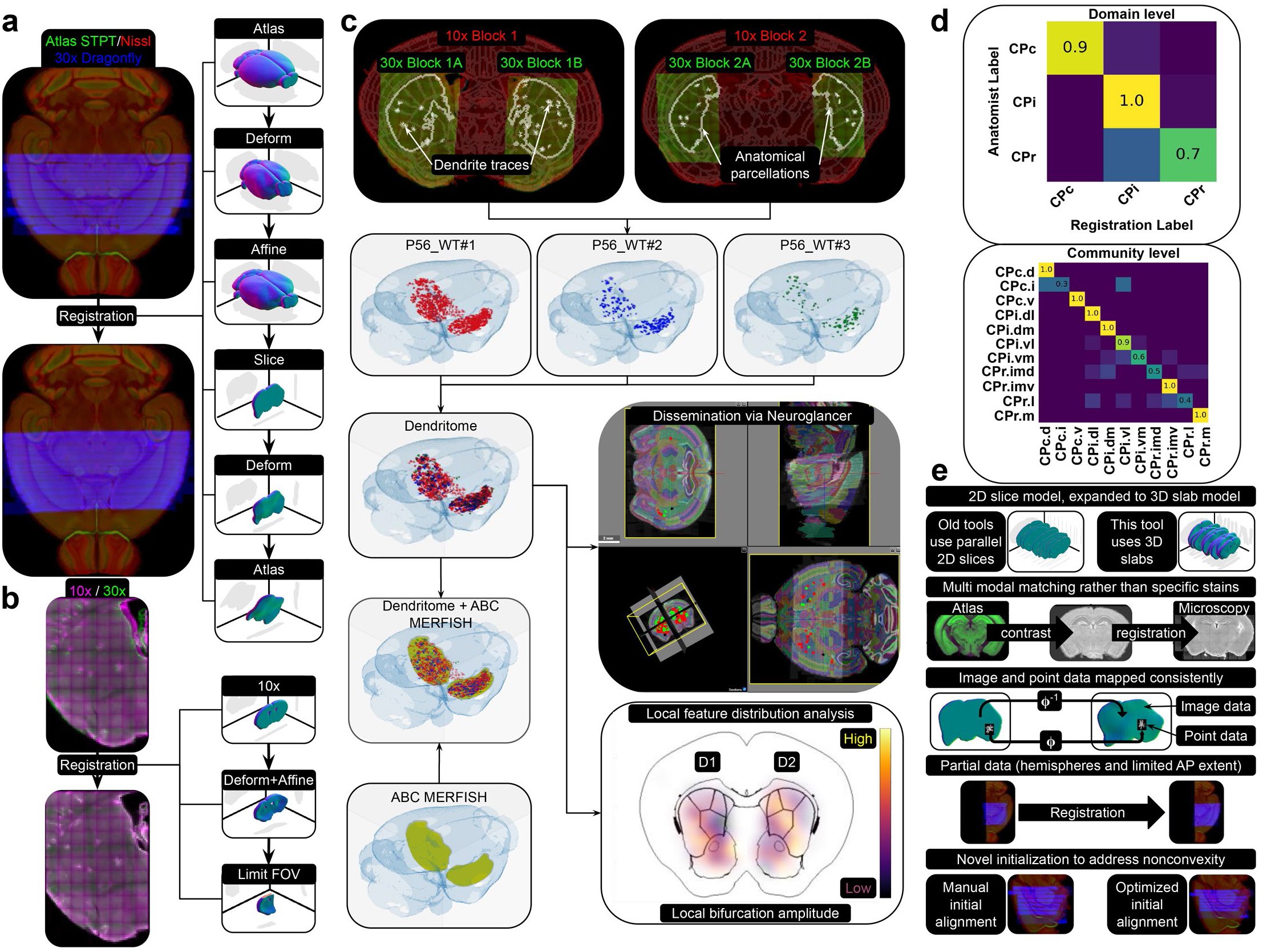
Overview of mapping images (10x atlas mapping, 10x to 30x ROI mapping) and neuron annotations (reconstructions) from thick-sectioned tissue into the CCF. **a,** The left shows a sequence of thick tissue sections (blue) for P56_WT#2 before and after registration to CCF (green/red). The right shows a sequence of transformations to iteratively estimate unknown parameters. **b,** The left shows a section imaged at 10x (magenta) aligned to the same tissue imaged at 30x. The right shows a series of transformations that iteratively estimate unknown parameters. **c,** The top shows two examples of 10x (red) and 30x (green) images registered together, with neuron reconstructions shown on the 30x image. The second row shows MSNs from different brains that are mapped into the CCF. The left panel demonstrates the joint analysis capability of the dendritome with other datasets, such as the Allen Brain Cell Atlas. Two downstream products are shown: 1) a 3D web-based visualization using Neuroglancer, and 2) local analysis of morphological feature distribution. **d,** Accuracy validation of registered anatomical regions compared to regions manually assigned by an expert neuroanatomist at the levels of 1) the rostral-intermediate-caudal axis and 2) corticostriatal axonal projection communities. **e,** Our registration algorithm introduces several key novel features: 1) handling 3D slabs instead of 2D slices, 2) accommodating multimodal images by estimating contrast differences, 3) ensuring inverse consistency of deformations in the LDDMM framework, 4) handling partial data, and 5) initializing nonconvex optimization using known anatomical targets for each thick tissue section.

Throughout this mostly automated pipeline, we implemented key manual quality control steps to ensure the high quality of the final outputs, e.g., eliminating neurons too close to the cut-surface and hence likely missing branches or those with low labeling signals (see Methods). Importantly, each final MSN reconstruction was manually reviewed by two experienced reviewers (about 5-10 minutes to manually correct each neuronal reconstruction), and each reviewed reconstruction was further validated by a third high-level validator. A final quality check was performed by expert scientists leveraging the software tools used by NeuroMorpho.Org^53^ to semi-automatically identify and correct the neuronal tracing SWC files^54,55^ for any structural irregularities (e.g., long connections, trifurcations, overlapping branches, etc.; Fig. 1a).

Together, this study generated a total of 3,762 fully reconstructed D1- and D2-MSNs from five P56 mouse brains, three 12-month old control, and four 12-month HD mouse brains (Fig. 1b). The latter is an HD knock-in mouse model (Q140 mice) expressing endogenous murine mutant Huntingtin (mHtt) with 140 CAG repeats^56,57^. Our P56 dataset, with 1,341 D1-MSNs and 1,125 D2-MSNs, distributed throughout the striatum of five brains, can be considered a high-quality WT MSN “dendritome” dataset. The 12m WT control and HD mouse brains, with a total of 617 D1-MSNs and 679 D2-MSNs, will allow us to examine the impacts of aging and mHtt on the MSN morphology. Compared to the historical archived data^9,58^, our study approximately doubles the number of MSN reconstructions compared to all of the previous studies of the rodent striatum (3761 MSNs in NeuroMorpho.Org v8.6), and with the advantage of more completed MSN dendrites (i.e. 500 μm thick sections in this study compared to 100-350 μm in previous studies). In terms of MSN types, prior studies cumulatively reconstructed only 219 and 290 identified D1- and D2-MSNs (NeuroMorpho.Org)^32,35,36,47^. Thus, our study provided a 4x-6x increase in the number of reconstructed D1- and D2-MSNs compared to the previous knowledge base.

### Multiscale registration of thick-sectioned tissue and mapped neuronal reconstructions to 3D brain reference atlas

Brainwide analysis of single-neuron morphology requires accurate registration of cell soma to the established mouse 3D brain reference atlas (i.e., CCFv3)^44,45^. To this end, we developed a novel deformable image registration algorithm that incorporates the Allen Atlas images^45^ with a recently developed set of segmentation labels in the dorsal striatum^59^. The program can register a sequence of 3D tissue slabs (i.e., 500μm) at 10x resolution, and to a series of one or more regions of interest (i.e., striatum) per slab at 30x resolution (Fig. 2a). An overview of our pipeline and its downstream goals are summarized in Fig. 2a-c. This generalizable approach enables the integration of our current datasets with other morphological data sets that are registered to the same 3D mouse brain reference space^44,45^ (Fig. 2c).

Our pipeline required several novel contributions (Fig. 2e). First, we used a sequence of 3D slabs, as opposed to a sequence of 2D parallel slices, and are able to represent “out-of-plane” deformations as well as oblique sectioning (see Methods). Second, we implemented multi-modal alignment, without requiring special stains to connect neighboring slices^60^. Third, we integrated the computational anatomy random orbit model^61^, allowing rigorous quantification of anatomical shape in a metric space, and a guarantee of well-posed inverse transformations for mapping image and point data consistently, as well as Jacobian factors for tracking changes in cell density. Fourth, unlike alternative registration approaches that are designed to be symmetric^62^, we support partial data, including image hemispheres, and fields of view that do not contain the whole brain (i.e., striatal regions of interest, or ROIs). Lastly, we used a novel approach for incorporating prior information into initial conditions, solving a low dimensional optimization problem to estimate initial anterior-posterior positioning and slab thickness (see Methods).

By composing the estimated transformations, we can map all neurons to the CCFv3 and assign them probable anatomical regions (Fig. 2d). Using 4 sections from 2 brains, we evaluated the concordance of these assignments to our previously established striatal anatomical divisions^63^ by a neuroanatomical expert. At the three striatal rostral-caudal levels, we achieve 91.2% agreement for neurons in the CPc, 100% agreement for neurons in the CPi, and about 75.5% agreement for those in CPr. The latter includes several neurons that our mapping method assigned to the CPi, while a neuroanatomist assigned them to CPr (Fig. 2d). At the further refined 11 striatal communities, which were established based on corticostriatal axonal projection patterns^63^ (Fig. 2d), we also observe a generally high concordance, but there are more disagreements for smaller structures.

### Integrative Analyses of MSN morphometry with 31 quantitative features

With each digitally reconstructed MSN in SWC file format, we computed 31 quantitative morphometric features (Supplementary Fig. 1 and Supplementary Table 1), 28 from L-Measure^50,51^ and 3 from TREES Boxtools^52^. These include 14 length related features, 7 angle related ones, and 10 branching complexity related ones. The length related L-Measure features measure total dendritic length (Length), the longest branches (Max_PathDistance, Max_EucDistance), and Euclidean lengths of individual dendritic compartments free of branchpoints (ABELs and BAPLs)^51^. The angle-related parameters include Bif_ampl_local, Bif_ampl_remote, Bif_tilt_local, Bif_tilt_remote, Bif_torque_local, Bif_torque_remote, which quantify various angle relationships between branches. The branching complexity related ones include bifurcations (N_bifs), number of dendritic ends (N_tips), and branches emanating from the soma (N_stems). Our study also uses three TREES Toolbox measures^52^ of dendritic spanning (convexity), centripetal bias (the degrees to which the dendritic branches are pointing towards the soma), and balancing factor (a graph-based measure of the balance between local dendritic wiring and path length to the soma). To our knowledge, no prior studies have utilized such extensive sets of morphometric features on a large fully reconstructed and validated mammalian MSN dataset, our study provides an opportunity for unbiased examination of the relationships among these quantitative readouts. We performed a hierarchical clustering based on the correlation of all the shape statistics in our P56 MSN datasets (Supplementary Fig. 2a). As expected, we found a large highly-correlated cluster of features including most length-related features (e.g., Max_EucDistance, Length, ABEL_all, Height, etc.) plus several branching related features (N_bifs, N_tips, N_stems), reflecting relatively symmetrical, spherical, and highly branched patterns of MSN dendrites. We also found the expected high correlation between centripetal bias and balancing factor (Cor=0.97), as strong centripetal bias corresponds to shorter path to the soma, which means larger balancing factor^64^. Our analysis also reveals novel relationships, such as the significant positive correlations between balancing factors and centripetal bias with Bif_tilt_remote (Cor=0.67 and 0.65 respectively) and negative correlations with Bif_ampl_remote (Cor=-0.79 and -0.62, respectively; Supplementary Fig. 2a). The former measures the angle between the previous node of the current bifurcating “parent” node and its “child nodes”; while the latter measures the angle between two bifurcation points and terminal tip or between two terminal points. This finding suggests the former but not latter measure is more closely correlated with the “root angle” (the angles of dendritic segments deviating from a direct path to the soma) used to calculate centripetal bias^64^. Another interesting and unexpected relationship is the strong negative correlation between “Terminal degrees” (total number of tips a segment will terminate into) and the sum of the product of contraction with Euclidean length and path length averaged on all the compartments, i.e.. ABEL_Terminal and BAPL_Terminal (i.e., -0.66 and -0.68). In sum, our dataset provides rich new knowledge on the relationships among the morphometric measures and can help derive a relatively non-redundant subset of measures (Supplementary Fig. 2b), based on clustering relationships, to perform more detailed morphological analysis of MSNs.

### Differential morphometry of D1- and D2-MSNs across striatal regions

To our knowledge, no prior studies have systematically examined the morphological variations of MSNs across the entire anatomical domain of the mammalian striatum. The published dendritic morphologies of a limited number of D1- and D2-MSNs are either restricted to the dorsal striatum^35,36^, CPr.dl^32^, or unspecified striatal region(s). We examined all 2248 3D reconstructed D1- and D2-MSNs from five P56 WT mouse brains for differential morphological analyses (Fig. 3b; Supplementary Table 2). Since these brains are registered to the CCFv3, our analyses can be carried out at increasingly refined anatomical resolutions from three striatal levels (CPr, CPi, CPc) to 11 striatal communities (CPr.m, CPr.imd, CPr.imv, CPr.l, CPi.dm, CPi.dl, CPi.vm, CPi.vl, CPc.d, CPc.i, CPc.v; Fig. 3a; Supplementary Table 3, Supplementary Table 4)^63^. At the community level, the number of MSNs range from 57 in CPr.imd to 344 MSNs in CPi.vm, mirroring the relative volumes of these striatal regions. Moreover, D1- and D2-MSNs are generally distributed equally among the reconstructed MSNs in each striatal community (Fig. 3b). Prior studies of D1- and D2-MSN morphologies, using only a few L-Measure readouts and/or Sholl analysis, have shown that D2-MSNs have shorter total dendritic length (Length), fewer primary branches (N_stems) and fewer total branch points than D1-MSNs (N_Branch)^35,36^.

**Fig. 3.**
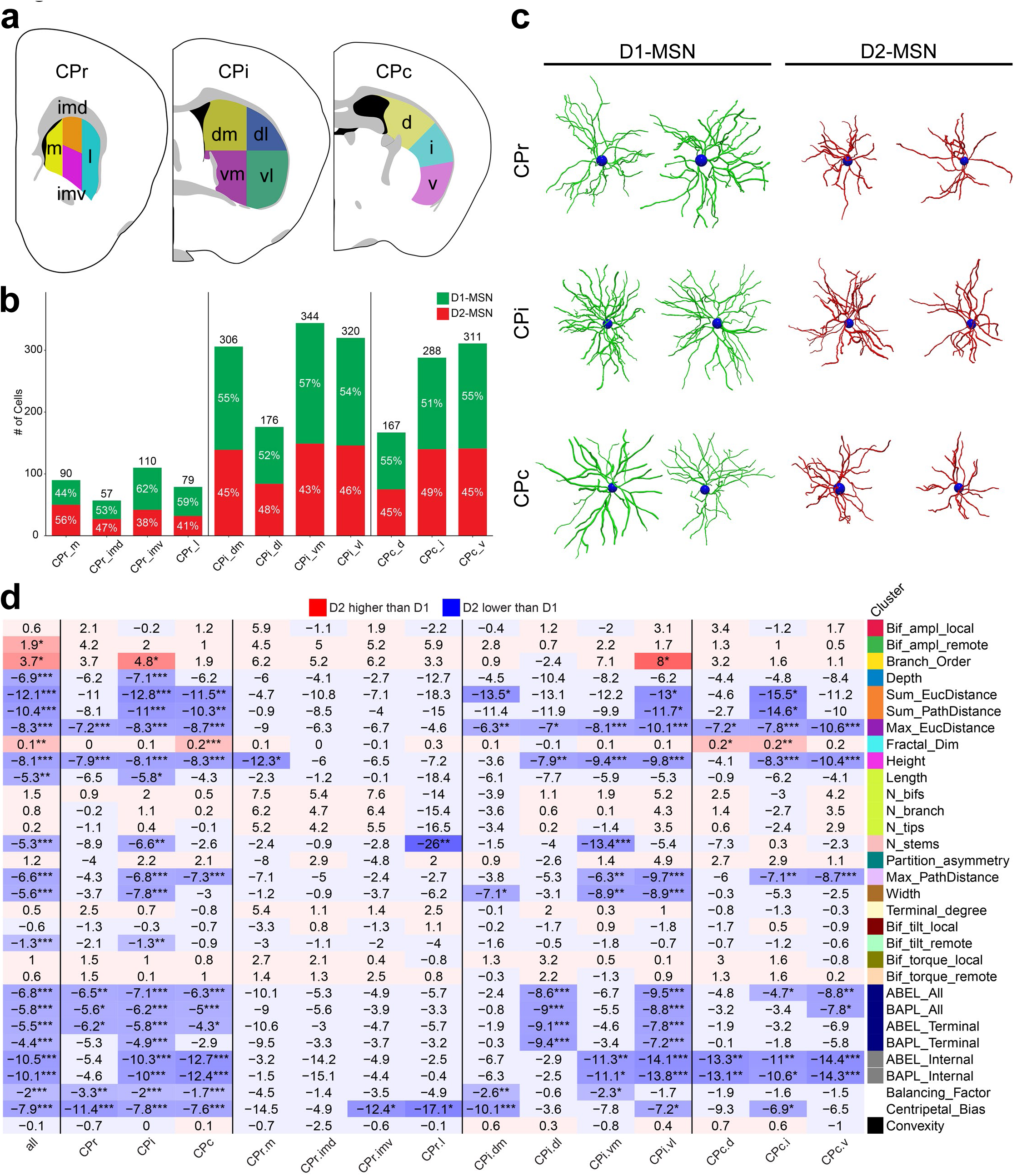
Morphological differences between D1- and D2-MSNs mapped to striatal domains. **a**, Division of the striatum to 11 striatal communities along the rostral-caudal axis (CPr, CPi, CPc) based on corticostriatal axonal projection patterns, including CPr.m, CPr.imd, CPr.imv, CPr.l in rostral CP, CPi.dm, CPi.dl, CPi.vm, CPi.vl in intermediate CP and CPc.d, CPc.i, CPc.v in caudal CP. **b**, The distribution of MSNs mapped to 11 striatal communities, and the distribution by D1- and D2-MSNs from P56 brains. **c,** Examples of reconstructions of D1- and D2- MSNs from rostral, intermediate and caudal CP. **d,** The heatmap of percentage changes of morphometric features of D2-MSNs compared to D1-MSNs, overall, in 3 striatal subregions and 11 striatal communities (two-sample t-test was performed). Red denotes morphometric features D2-MSNs higher than D1-MSNs, and blue denotes morphometric features D2-MSNs lower than D2-MSNs. The color bar denotes the grouping of morphometric features. Caudate Putamen (CP) Abbreviations: CPr, CP rostral level; CPi, CP intermediate level; CPc, CP caudal level; CPr.m, medial community of CPr; CPr.imd, intermediate dorsal community of CPr; CPr.imv, intermediate ventral community of CPr; CPr.l, lateral community of CPr; CPi.dm dorsomedial community of CPi; CPi.vm, ventromedial community of CPi; CPi.dl, dorsolateral community of CPi; CPi.vl, ventrolateral community of CPi; CPc.d, dorsal community of CPc; CPc.i, intermediate community of CPc; CPc.v, ventral community of CPc.

Our study broadly extends the D1- and D2-MSN morphological comparison to the entire striatum with 31 morphometric statistics (Supplementary Table 2). First, at the whole striatum level, 21 of the 31 morphometric features significantly differ between D1- and D2-MSNs (Fig. 3d). The significant features are more prominent in the CPi and CPc levels than in CPr, which could partially due to the different number of reconstructed MSNs per region (Fig. 3b-d). At the community level, the D1- and D2-MSNs appear to differ the most in the seven CPi and CPc communities, with the most consistent differences being that D1-MSNs have longer Max_EucDistance than D2-MSNs (Fig. 3d). Other length related features that are longer in D1-MSNs include individual branch segments (ABELs and BAPLs) and Max_EucDistance. Importantly, although the number of primary dendrites (N_stems) is fewer in D2-MSNs than D1- MSNs, consistent with a prior study^36^, other branching related features (N_bifs and N_tips) are not significantly different between the two MSN types (Fig. 3d). One novel finding here is that D2- MSNs consistently have a higher Fractal Dimension, which quantifies the fractal-like behaviors of neuronal branching, with higher Fractal_Dim having more contributions of fine-scale branches relative to coarse ones^65^.

At the community resolution, multiple D1- and D2-MSN differences do not appear to be attributed to MSN numbers, but instead are likely to be biological (Fig. 3b, d). In the CPi, the two lateral communities have significant D1/D2-MSN differences in ABEL-related features while the two medial communities have no differences, despite the fact that the CPi.dl has almost half the number of MSNs compared to the other communities (Fig. 3d). Similarly, D2-MSNs in CPi.vl have higher Branch_Order (i.e. order of branches with respect to the soma) as compared to D1- MSNs (Fig. 3d). Additionally, a few communities in CPr (CPr.imv, CPr.l), CPi (CPi.dm, CPi.vl), and CPc (CPc.i) show interesting D2-MSN branching patterns such that their trees are less straight, and the tips of their dendritic arbors point more towards soma in comparison to D1-MSN (Fig. 3d). Meanwhile, all the striatal communities of about 300 MSNs, except CPi.dm, show significant D1/D2-differences in Height, Max_PathDistance, ABEL_Internal, BAPL_Internal. Together, our analyses provide a nuanced view of the D1- and D2-MSN morphological differences and emphasize the importance of examining the MSN types in precise striatal anatomical regions.

### Striatal region-specific morphological features shared by D1- and D2-MSNs

Our analysis thus far showed that the D1- vs D2-MSN morphological differences are variable across striatal levels and communities, with some regions and features more robustly differing than others. We next hypothesized that perhaps the MSN morphological differences across striatal regions could be more consistent. We first visualized the mean values of every morphometric parameter for MSNs at the community levels, each normalized by the median value of the parameter in the brain (Fig. 4a). The MSNs at the CPr levels tend to have lower total length and branching related parameters than the median values of the whole striatum, but interestingly longer individual dendritic segments (ABELs and BAPLs) and bifurcation remote angles (Bif_ampl_remote) than the median values (Fig. 4a). The MSNs in the CPi and CPc communities, in contrast, tend to have higher total dendritic length and more branches. However, individual communities have their own specific features, e.g., the CPi.vm has higher ABEL values while the CPi.dm and CPc.v do not seem to have higher dendritic length or number of branches.

**Fig. 4.**
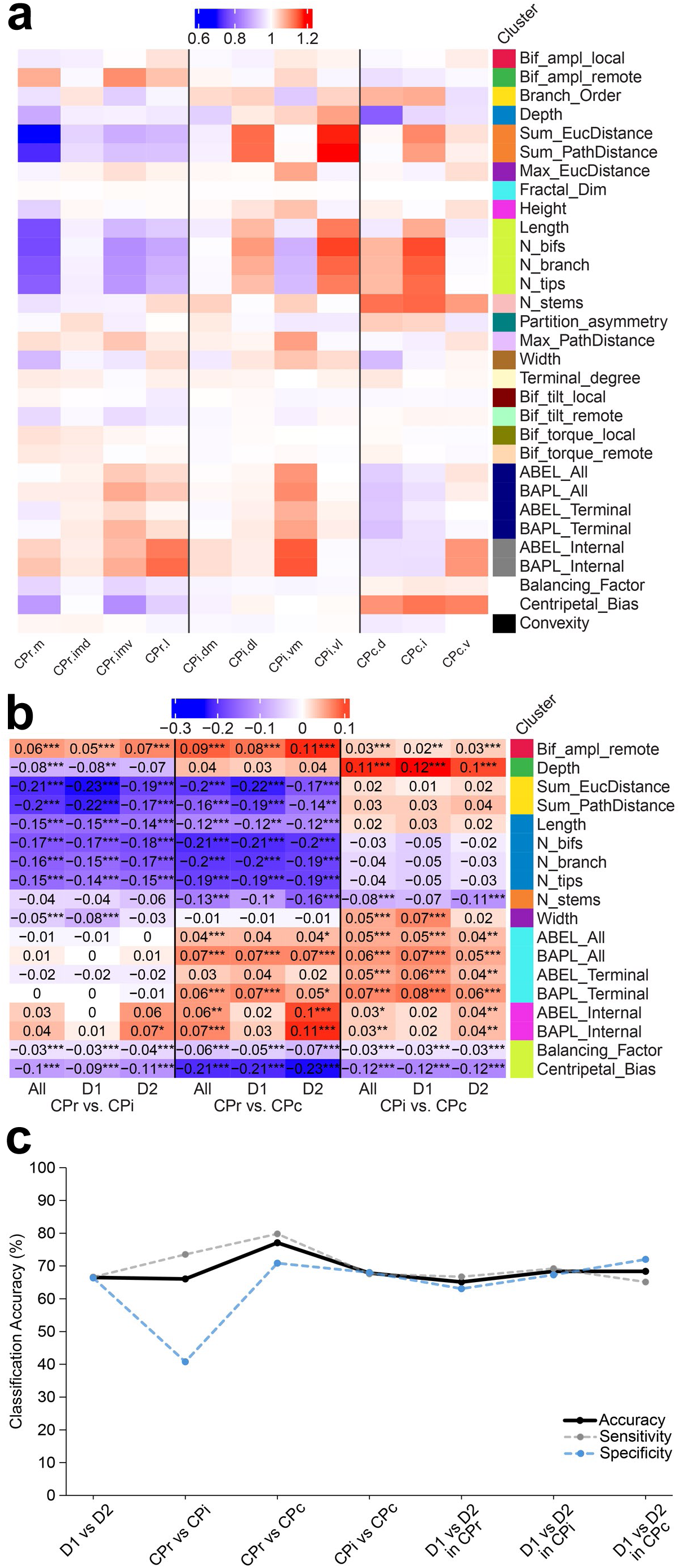
Regional distribution and regional differences of MSN morphology. **a**, The distribution of median-normalized morphometrics of MSNs across 11 striatal communities. Blue denotes values are lower than the brain median and red denotes values are higher than the brain median. **b**, The pairwise regional comparison (CPr vs. CPi, CPr vs. CPc, and CPi vs CPc) of median- normalized morphometric features for all MSNs, D1-, and D2- MSNs (two-sample t-test); only morphometrics with statistically significant regional differences are shown. **c**, Supervised classification results of the separation of D1- and D2- MSNs and the separation of CPr, CPi, and CPc, using Support Vector Machine (SVM) classification.

We next compared the differences in MSN morphometry, either combined or separately by D1 or D2-MSN only, across striatal levels (Fig. 4b). This analysis reveals remarkably consistent and significant differences in 18 morphometric measures, 3 total length related features, 4 branching features, 6 ABEL/BAPL features, and balancing factor as well as centripetal bias (Fig. 4b). Importantly, CPr MSNs, regardless of D1 or D2, are smaller in length and branching complexity but comparable in ABEL_all/ABEL_terminal compared to the CPi or CPc MSNs. Comparing CPi and CPc MSNs, regardless of D1 or D2, show similar total length and branching related features but differ significantly in all 6 individual dendritic branch lengths (ABELs/BAPLs), which are higher in the CPi (Fig. 4b). Finally, both balancing factor and centripetal bias show a significant and graded increase from CPr to CPi and CPc, and an inverse of this trend is seen in the bifurcation remote angles (Bif_ampl_remote; Fig. 4b). Overall, the morphometric differences across striatal regions described here remain generally consistent across individual brains (Supplementary Fig. 3, 4), suggesting the robustness of such findings and their potential utility as reference MSN morphological variations in the WT striatum.

We next asked whether support vector classification (SVM) models^66^ could be trained to utilize morphometric features to classify striatal MSN D1/D2-types and/or subregions. The test sets grouped by MSN types or striatal level all showed similar prediction accuracy (greater than 65%), with classification of CPr vs CPc particularly accurate (Fig. 4c; Supplementary Table 5). Thus, based on the existing striatal anatomical annotation, we found robust and consistent differences in dendritic morphometry, regardless of D1- and D2-MSNs, across the three striatal rostral-caudal levels.

### Unbiased discovery of MSN dendritic morphological modules based on voxelated “striatal boxes” in the CCFv3 space

Since striatal anatomical locations seem to have a reproducible influence on the MSN morphometry (Fig. 3d; Fig 4a, 4b; Supplementary Fig. 5), we hypothesized that an analysis of such striatal subregion effects on MSN morphology at a finer granularity could provide deeper insights into MSN dendritome organization. To this end, we understand the limitations of subdividing striatal domains based on known anatomical landmarks only (i.e., the 3 striatal levels) or based on the corticostriatal projectome (i.e., 11 communities)^63^. Although the latter is registered to the CCFv3^59^ (Supplementary Table 4) and used in our study, we recognize the community-level registration is an approximation as we do not have a simple experimental way to discern the specific corticostriatal axonal pathways to validate such community assignments. To overcome these challenges, we decided to pursue a voxel-based approach by dividing the entire CCFv3 into 7020 cubic boxes, each box being 500 µm per side. In this reference voxelated CCFv3 space, there are 210 boxes encompassing the entire striatum. We next analyzed MSN morphological variations at the level of boxes, an approach we termed “box-based morphometric analysis” (Fig. 5a). Briefly, all the reconstructed MSNs were mapped to individual striatal boxes, with each box having a fairly equal mixture of D1- and D2-MSNs, as expected (Fig. 5b; Supplementary Fig. 6a-6c). Next, we calculated one representative value for each of the 21 non-redundant morphometric features for the MSNs in a given box (see Methods) and used the resulting values as a summary morphometric descriptor for the MSNs in each box (a feature we referred to as “eigen-morphometry” or “eigen- morph”; Fig. 5a). This approach is inspired by the omics analysis algorithm WGCNA^38^, as ours is similar to WGCNA since we use the first principal component of each morphometric feature for all MSNs in a box to generate the eigen-morph values. We next performed hierarchical clustering of eigen-morphs representing 210 non-empty striatal boxes, which reveals a robust cluster dendrogram (Fig. 5b). Based on this clustering dendrogram, we selected 7 clusters as MSN “dendritic modules (DM)” (i.e. DM1-DM7). To identify the likely distinguishing morphometric features of these modules, we visualized the features of the top 25 representative cells for each of the dendritic modules, with cells ranked based on their loadings in their respective DM modules (Fig. 5c). Importantly, each of the dendritic module representative MSNs show characteristic co- variations across the 21 morphometric features, suggesting that similarities in a distinct subset of morphometric features are driving the dendritic module formation, again a phenomenon analogous to the identification of gene coexpression modules with WGCNA^38^.

**Fig. 5.**
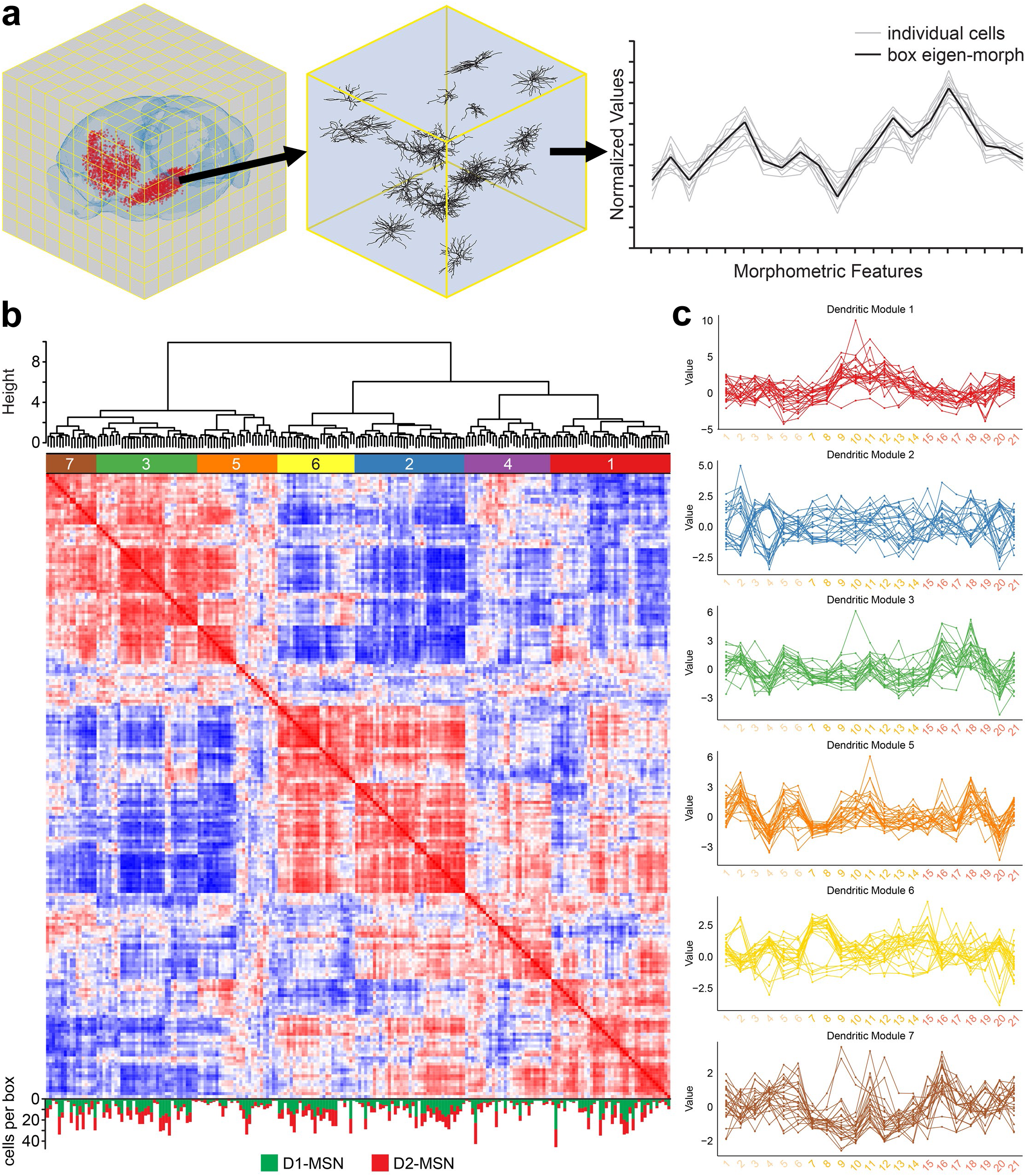
Unbiased box-based analysis of MSN morphologies in the whole striatum. **a**, The division of the CCF-registered mouse brain into equal-sized (500μm) cubic boxes (left). MSNs were allocated into corresponding boxes based on mapped spatial locations (middle) and eigen- morph were extracted to represent the overall morphological variations within each box (right). **b**, The hierarchical clustering of boxes based on box eigen-morph, including the dendrogram of the box clustering, the pairwise box similarities (red, more similar between box eigen-morph; blue, less similar between box eigen-morph), and D1- and D2- MSNs distribution per box. **c**, The morphological variations of 21 non-redundant morphological features of the representative neurons in each morphological territory (MT).

We next examined the distribution of normalized morphometrics (see Methods; Supplementary Fig. 7) across all boxes, grouped by DM modules (Fig. 6a). Each module shows distinct combinations of morphometric characteristics. MSNs in DM3, DM5 and DM7 generally exhibit smaller total length, and fewer branches, and average branch length and centripetal bias. Two other modules, DM1 and DM4 also have shorter dendritic length-related features (e.g. Lengths, Max_EucDistance), smaller bifurcation angles, and fewer branches; while they are distinguished from DM3, DM5, and DM7 due to longer ABEL-related features. Lastly, neurons in DM2 and DM6 have higher dendritic length and branching complexity but shorter individual dendritic branches. Representative MSNs from 6 of the DM modules, selected from those with high contribution (loading) to the respective module, are shown in Fig. 6b, further illustrating the unbiased dendritic morphological module-based morphological variations in the striatum. Moreover, by configuring our imaging and reconstruction datasets for use with the Neuroglancer framework, we provided a web-based, interactive visualization of the top 25 representative MSNs from these DM modules in the 3D CCFv3 space (Fig. 6c).

**Fig. 6.**
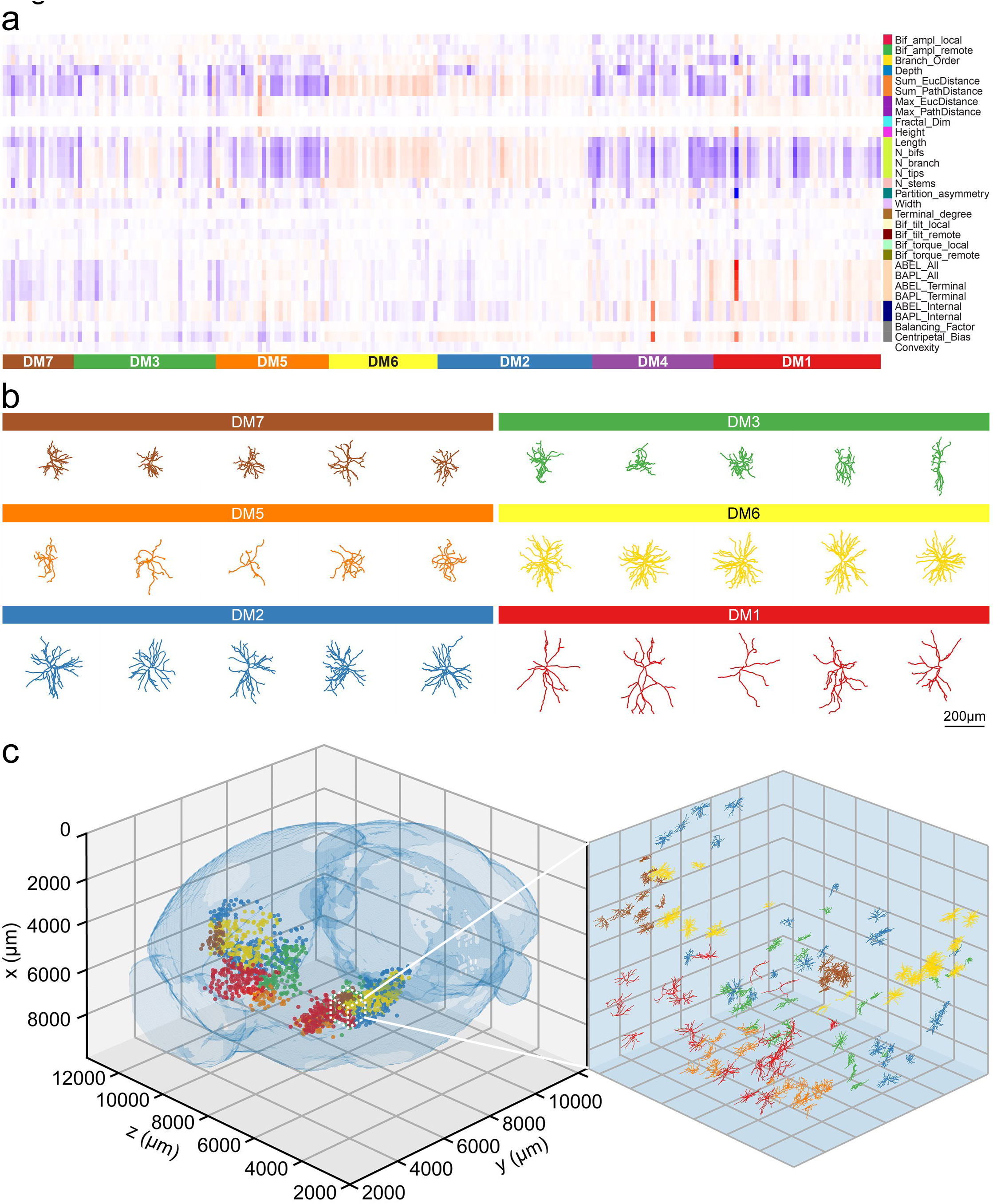
Morphological distribution of MSNs in dendritic modules (DMs) and spatial variations of MSN morphologies in the DMs. **a**, The distribution of median-normalized morphometric features of MSNs, ordered and grouped by DMs (red: value above the brain median, blue: value below the brain median). **b**, The morphology of representative MSNs in each morphological territory (MT), showing distinct morphological variances. **c**, The spatial distribution of MSNs in the identified MTs.

### MSN dendritic morphological territories receive distinct corticostriatal inputs

We next asked whether spatial distribution of the boxes within the same dendritic morphological module is random or not, with a focus on the distribution of boxes within the same module that are physically interconnected in CCFv3 space. Importantly, we found that the distributions of the connected boxes in the DM modules are statistically significant from non-random, spatially connected distributions (Fig. 7a). There are 6 DMs containing connected box cluster with ≥7 boxes: 25 boxes in DM1, 26 in DM2, 15 in DM3, 9 in DM5, 19 in DM6, and 7 in DM7 (Fig. 7a; Supplementary Fig. 6d, e). DM4 had boxes fairly evenly distributed throughout the striatum with no more than five boxes connected and most in singles or pairs. We refer to these connected boxes (≥7 boxes) for DM modules as MSN morphological territories (MT), e.g. MT-1 for DM1, MT-2 for DM2, MT-3 for DM3, MT-5 for DM5, MT-6 for DM6, and MT-7 for DM7. These MSN MTs are identified without incorporating prior anatomical annotations and most of them span several registered striatal levels or communities in CCFv3 (Fig. 7b, 7c). As an illustration of the robust differences in MSN dendritic morphological features in these MTs, trained SVM classifier can categorize individual MTs against each other with moderate to high prediction accuracy (63-87%; Fig. 7d; Supplementary Table 5).

**Fig. 7.**
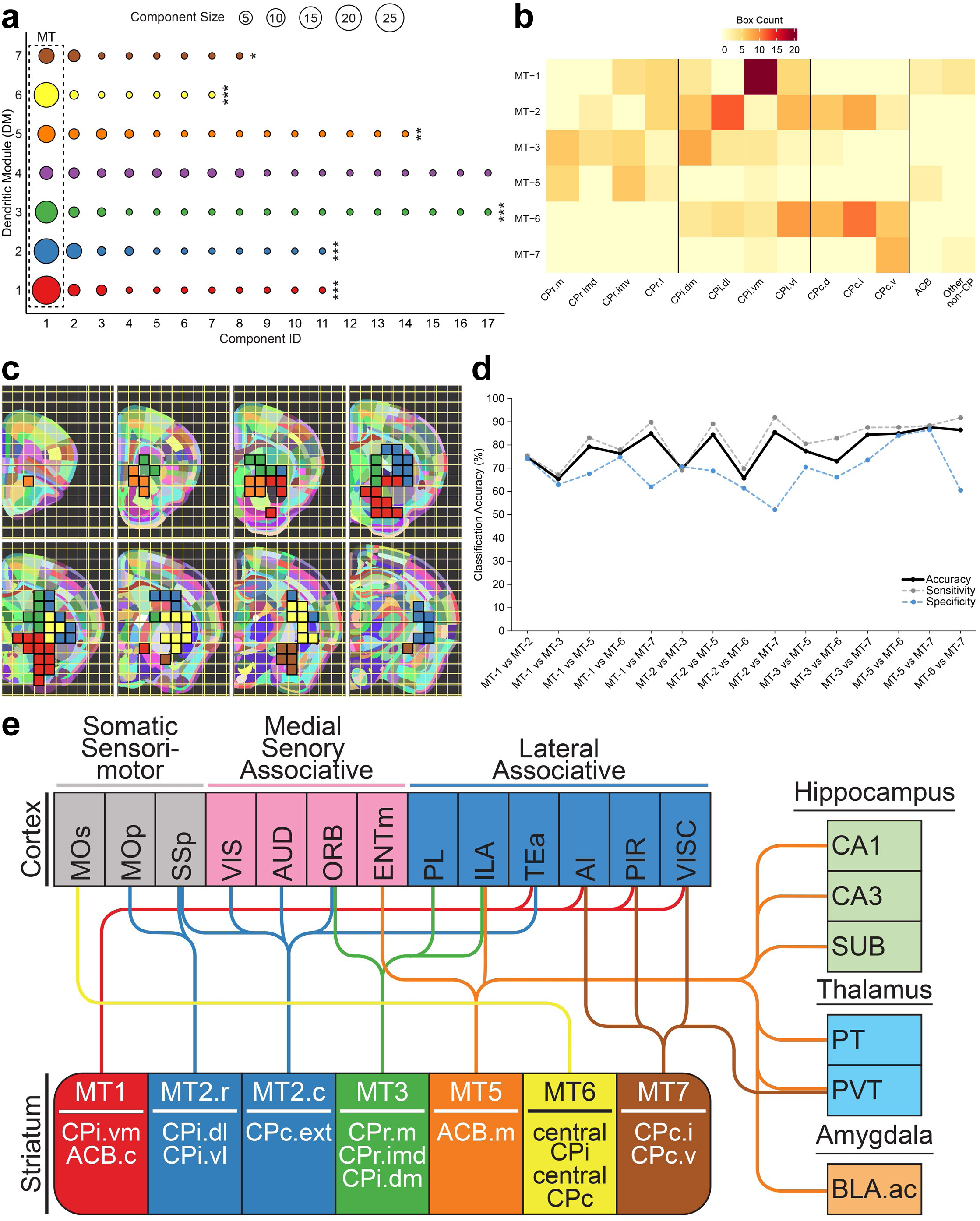
Identification, spatial distribution, and cortical inputs to striatal morphological territories (MTs) derived from connected DMs. **a**, The distribution of the sizes of connected components within each DM to define the components with >= 7 boxes as MTs (chi-square goodness-of-fit test). **b**, The distribution of MT-associated boxes across striatal communities. **c**, The spatial distribution of the boxes in MTs, from rostral to caudal CP, in the CCF-registered mouse brain. **d,** The supervised classification results of separating 6 MTs, using Support Vector Machine (SVM). **e,** Converging cortical and subcortical inputs into striatal MTs. Cortical abbreviations: AI, agranular insular area; AUD, auditory area; ENTm, entorhinal area, medial; ILA, infralimbic area; MOp, primary motor area; MOs, secondary motor area; ORB, orbitofrontal area; PIR, piriform cortex; PL, prelimbic area; ; SSp, primary somatosensory area; TEa, temporal association area; VIS, visual area; VISC, visceral area. Subcortical abbreviations: CA1, hippocampal formation, field CA1; CA3, hippocampal formation, field CA3; SUB, subiculum; PT, paratenial nucleus of the thalamus; PVT, paraventricular nucleus of the thalamus; BLA.ac, basolateral amygdalar nucleus, anterior, caudal domain.

We next asked whether each MT could receive a consistent set of cortical inputs that overlaps with known functional domains in the striatum (Fig. 7e)^63^.

MT-1 (DM1) is present in a contiguous region that spans the posterior wing of the nucleus accumbens (ACB) at approximately ARA 48-50 and the adjacent ventromedial striatum including the CPr.l.vm, CPi.vm.vm, and CPi.vm.v. These are limbic regions in classic models of the basal ganglia^67^. The MT-1 region integrates cortical inputs from perirhinal, temporal association, visceral, and all agranular insular areas, as well as piriform cortex, among others.

MT-2 (DM2) is the only cluster to have two non-contiguous sectors, one more rostral and one more caudal. The more rostral group lies along the lateral edge of the caudoputamen (striatum) at its intermediate (commissural) level, corresponding to the lateral zones of CPi.dl.d, CPi.dl.imd, CPi.vl.imv, CPi.vl.v, and CPi.vl.vt. These domains constitute the body homunculus representation, and integrate primary somatomotor and primary somatosensory information for each body region. The more caudal group lies in the caudal extreme of the caudoputamen (CPc.ext), also known as the tail region. This domain integrates polysensory inputs from auditory, visual, and somatosensory cortices, along with higher-order temporal and orbitofrontal cortical regions which themselves integrate multiple sensory inputs^63,68^.

MT-3 (DM3) primarily occupies domains of the dorsomedial caudoputamen at the rostral (CPr.m) and intermediate levels (CPi.dm.d, CPi.dm.im, CPi.dm.cd, CPi.dm.dl) with some ventral encroachment into the adjacent ventromedial caudoputamen (CPr.imv, CPi.vm.vm, CPc.d.vm). Although these striatal domains each receive a unique complement of cortical and amygdalar inputs, collectively they are innervated by the prelimbic (PL), orbital (ORB), infralimbic (ILA), anterior cingulate, auditory, ectorhinal, posterior parietal association, retrosplenial, temporal association, and visual cortical areas, and basomedial and basolateral amygdala^63,69^. These inputs innervate more than one domain in this group of domains, meaning this region is enriched in these inputs, and in particular the medial prefrontal areas PL, ORB, and ILA provide the most innervation to this region. The CPr.m region covered by MT-3 contains domains that integrate cortical inputs related to complex cognitive and associative functions, with likely roles in Pavlovian fear conditioning, saccadic oculomotor control, high-order polysensory integration, and visuospatial information processing for orientation and navigation^63^.

MT-6 (DM6) occupies the central region of the caudoputamen at its intermediate level, extending caudally to the middle region of the caudal caudoputamen. This corresponds, at the intermediate level, to the medial zones of domains CPi.dl.d, CPi.dl.imd, CPi.vl.imv, CPi.vl.v, and CPi.vl.vt and CPi.vl.cvl. As described above, these domains integrate somatomotor and somatosensory cortical inputs, forming the body homunculus representation in the caudoputamen. Laterally within these domains, there is a greater density of primary motor and primary somatosensory inputs, which rostral MT-2 occupies; but medially within these domains, there is a greater density of secondary motor cortex (MOs) inputs for each of the body regions, which MT- 6 is positioned to receive. Separate basal ganglia pathways for primary and secondary motor information have also been described in primate^70^. In caudal caudoputamen, MT-6 is situated in the middle of the nucleus where the MOs inputs terminate densely.

Finally, MT-7 (DM7)) overlaps with ventromedial CPc (part of CPc.v.vm; Fig. 7b, c). This is the only dendritic morphological territory that is confined to one striatal community. Interestingly, this region is known to integrate inputs from visceral, agranular insular, and piriform cortex, which are known to process visceral and inner state signals and regulate autonomic functions^63^.

Together, our unbiased analysis of the striatal MSN dendritome reveals a novel organization of the murine striatum based on dendritic morphological territories, which receive functionally distinct cortical inputs.

### Impacts of aging and the Huntington’s disease mutation on striatal D1- and D2-MSN dendritome

Neuronal dendrites are a site of active plasticity^71^, and are known to remodel during aging^72,73^, stress^31^, or in neurodegenerative diseases^74^. However, the brainwide morphological mapping approach has never been applied to study the dendritic remodeling of genetically defined neuronal types in normal aging or a disease condition. As a proof-of-concept, we imaged and reconstructed 548 genetically identified D1- and D2-MSNs from three 12m wildtype striata, and 748 D1- and D2-MSNs from four 12m Q140 mutant huntingtin (mHtt) knock-in mice^57^ (Figure 8a). The latter is an HD mouse model that shows progressive and selective pathogenesis restricted to the HD-vulnerable MSNs and certain cortical pyramidal neurons and is also an excellent model system to study HD GWAS modifier genes on selective neuronal pathogenesis^56,75^.

**Fig. 8.**
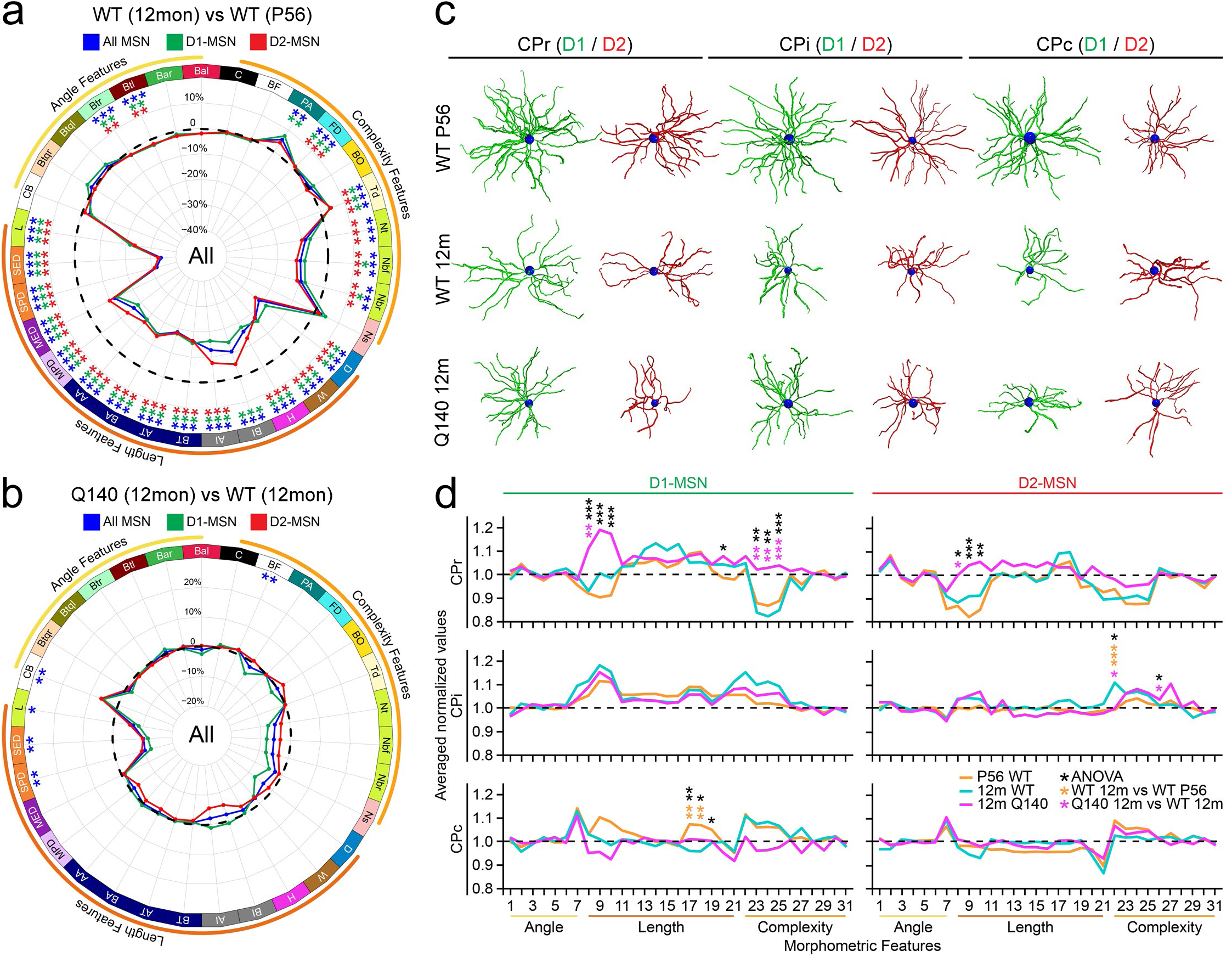
Morphometric differences between 12m WT and P56 WT MSNs and 12m HD and 12m WT MSNs. **a**, The percentage changes of morphometric features of MSNs from 12m WT (n=3) compared to P56 WT mice (n=5) across the entire striatum (two-sample t-test). **b**, The percentage changes of morphometric features of MSNs from 12m Q140 (n=4) compared to 12m WT mice (n=3) across the entire striatum (two-sample t-test). **c**, Examples of D1- (green) and D2-MSN (red) neuron morphology in CPr, CPi and CPc striatal levels from P56 WT, 12m WT and 12m Q140 brains. **d,** The distribution of the averaged, median-normalized morphometrics of MSNs from P56 WT (n=4), 12m WT (n=3) and 12m Q140 mice (n=4) in CPr, CPi, and CPc, separated by D1- and D2-MSNs (one-way ANOVA, followed by Tukey’s HSD post-hoc analysis on 12m Q140 vs. 12m WT and 12m Q140 vs. P56 WT). *P < 0.05, **P < 0.01, ***P < 0.001; in **a** and **b**, blue asterisks for both D1- and D2-MSNs, green asterisks for D1-MSNs only, and red asterisks for D2-MSNs only; in **d**, black asterisks for ANOVA, orange asterisks for 12m WT vs P56 WT, and pink asterisks for 12m Q140 vs 12m WT from the post-hoc tests. Morphometrics abbreviations: 1) Bal, Bif_ampl_local; 2) Bar, Bif_ampl_remote; 3) Btl, Bif_tilt_local; 4) Btr, Bif_tilt_remote; 5) Btql, Bif_torque_local; 6) Btqr, Bif_torque_remote; 7) CB, Centripetal_Bias; 8) L, Length; 9) SED, Sum_EucDistance; 10) SPD, Sum_PathDistance; 11) MED, Max_EucDistance; 12) MPD, Max_PathDistance; 13) AA, ABEL_All; 14) BA, BAPL_All; 15) AT, ABEL_Terminal; 16) BT, BAPL_Terminal; 17) AI, ABEL_Internal; 18) BI, BAPL_Internal; 19) H, Height; 20) W, Width; 21) D, Depth; 22) Ns, N_stems; 23) Nbr, N_branch; 24) Nbf, N_bifs; 25) Nt, N_tips; 26) Td, Terminal_degree; 27) BO, Branch_Order; 28) FD, Fractal_Dim; 29) PA, Partition_asymmetry; 30) BF, Balancing_Factor; 31) C, Convexity.

In 12m WT striata, D1- and D2-MSNs still exhibit significant differential morphological features in different striatal levels and relatively robust, MSN-type-independent morphological differences across striatal levels (Supplementary Fig. 8, 9, 10), preserving key features in P56 mice (Fig. 3d, 4a; Supplementary Fig. 3, 4). Comparing 12m to 2m MSNs, we found that both D1- and D2-MSNs exhibit a highly robust reduction of 13 dendritic length related features (e.g., Lengths, Max_EucDistance, ABELs) in all three striatal levels, while the CPc, but not CPr and CPi regions, also show a highly significant reduction of dendritic branching (Fig. 8a, 8c; Supplementary Fig. 10a). However, angle related features are relatively preserved. Together, these results suggest that aging led to a relatively uniform atrophy of both D1- and D2-MSN dendrites in terms of their length, and more striatal regional-selective reduction of dendritic complexity. At 12m, when comparing Q140 to WT MSNs, we found 5 features that significantly differ at the whole striatum level. HD MSNs are significantly shorter in dendritic length measures (Length, Sum_EucDistance, Sum_PathDistance) while larger in balancing factor and centripetal bias (Fig. 8b, 8c; Supplementary Fig. 10b).

We next analyze separately D1- and D2-MSNs to examine the aging as well as mHtt-induced effects on the distribution of mean morphometric features in three striatal levels relative to the median values in the whole brain. Interestingly, we found in D1-MSNs in the CPr region, WT brains at P56 and 12m show similar below median values in 3 length (Length, Sum_EucDistance, Sum_PathDistance) and 3 complexity (N_branch, N_bif, N_tips) related features, but Q140 HD MSNs show these features that are markedly above or comparable to the brainwide median values in the D1-MSNs (Fig. 8d). The Q140 vs WT at 12m are statistically significant in post-hoc analyses for these morphometric features in D1-MSNs, and Tukey’s HSD post-hoc analyses show significant differences between Q140 and WT at 12m in 4 out of 6 morphometric features. Similar but milder differences in the three lengths (but not complexity) related features also exist for D2-MSNs in CPr region between Q140 and WT at 12m, with one such feature being significant in post-hoc analyses (Fig. 8d). A few other significant differences in complexity features can be seen between Q140 and WT in D2-MSNs in CPi as well (Fig. 8d). Taken together, our study suggests that large-scale dendritome mapping can help sensitively reveal both aging and HD-associated MSN pathology that are selective to MSN types, striatal subregions, and distinct morphometric features.

## DISCUSSION

Currently, the molecular properties of single neurons, such as transcriptomes and epigenomes, can be profiled at large-scale across species^3,5,75,76,77^. However, single neuron properties beyond molecular signatures, such as morphology, electrophysiology and connectivity, may carry orthogonal information and help to more precisely define neuronal cell types^79,80,12,15^. To this end, our study demonstrates a novel integrated pipeline for the brainwide mapping of neuronal dendrites (i.e. dendritome mapping) for genetically-defined single neurons, using genetic tools, equipment and software that can be readily disseminated to the broader neuroscience community. Our pipeline for mapping neuronal dendrites has several advantages compare to other recent large-scale single-neuron morphological studies^11,12,13,14,15,16,79^. First, MORF3/Cre based genetic sparse labeling avoids the limitation of viral-based sparse labeling, e.g. labor intensiveness of manual injections, focal study of brain cell morphology, potential viral toxicities, and inability to study embryonic brains. Second, our tissue-clearing and imaging setups are commercially available and avoid using specialized imaging equipment (e.g., fMOST). Third, our semi-automated, novel dendrite reconstruction pipeline requires about 10-15 minutes of manual correction of reconstruction per neuron, compared to much longer time for other neuronal reconstruction^11^. Dendrites have a distinct function in neurocomputation and are an integral aspect of neuronal classification^81,82^, and dendritic analysis is more amenable in terms of large-scale studies compared to electrophysiology and single-neuronal axonal morphological studies. Thus, our study strongly argues for dendritome mapping as a scalable, single-neuron, brainwide morphological profiling approach that can be adapted by many labs to study diverse biological and pathological questions for mammalian neurons in vivo.

Our study provides the first striatum-wide D1- and D2-MSN dendritic morphological atlas in the adult mouse brain. Our study reveals shared length-related morphological differences between D1- and D2-MSNs across all striatal levels, replicating prior findings^35,36^, but also reveals novel findings such as differences in dendritic branch length (ABELs), balancing factors, and centripetal bias. At the community levels, there appears to be substantial variations in D1/D2-MSN morphological differences, even in communities with a similar number of reconstructed MSNs. These findings suggest one cannot generalize findings of D1/D2-MSNs from one striatal subregion to others.

An important advance in this study is our unbiased discovery of striatal MSN dendritic morphological territories, relying purely on the dendritic morphologies of CCF-mapped D1/D2- MSNs. Prior striatal domains relied on actual anatomical landmarks (e.g., rostral-caudal CP levels), corticostriatal axonal terminals (e.g., communities and domains)^63^, or molecular markers (e.g., striosome and matrix)^83^. Moreover, the morphological variations of D1- and D2-MSNs throughout the striatum domains were unknown prior to this study. Other recent large-scale studies of single-neuron dendrites and axons relied on prior anatomical knowledge^10,13,84^ or “microenvironments” (i.e., aggregate dendritic features of nearby partially reconstructed neurons)^85^, but such approaches are not suitable for brainwide analysis of the complete morphology of genetically defined neurons at fine resolution and without any prior anatomical constraints. To address such a question, we devised a generalizable strategy to perform brainwide unbiased analysis of the fully reconstructed dendritic morphology of genetically-defined single neurons. The core concept of this approach is to divide the whole CCFv3 brain into pixelated cubes of 500µm per side (i.e. boxes), and to analyze all the genetically-defined neurons mapped to each box to study their shared morphometry. Our design enables ready joint analysis (e.g. combining registered MSNs from different brains into the same box) or ready comparison across conditions (e.g. MSNs from different ages, disease status, or future studies by other labs). Our development of a summary statistic (eigen-morph), using WGCNA code^38^ to calculate the first principal component of non-redundant morphometric statistics for all the neurons within a box, enables us to perform striatum-wide clustering to identify modules of MSNs with similar morphological patterns. This unbiased CCFv3 box-based dendritic eigen-morph analysis should be readily applied to any future studies of spatial organization of genetically-defined neuronal dendrites throughout the brain, as long as the reconstructed single neurons are sufficiently dense enough to populate the boxes in brain regions containing such neurons.

Our study generates important new questions on the functional roles of dendritic morphological territories, and the possible mechanisms underlying such territories. An important clue to answering these questions is the distinct cortical inputs to the separate MTs, which appears to be broadly consistent with known corticostriatal circuit functions, e.g., separate primary and secondary motor cortices inputs to MT-2 and MT-5, and distinct visceral inputs to MT-1 and MT-7. Future studies are needed to examine whether cortical inputs are indeed causing the dendritic morphological variations in both D1- and D2-MSNs in these MTs, or if there are cell-autonomous determinants setting up such dendritic morphological patterns. Given that striatal MSN morphology is known to be highly plastic and responsive to experiences such as stress^31^ or dopamine signaling^28,30^, it is tempting to propose that circuit-specific neuronal communications could be a prime source of dendritic variations in MSNs and possibly for other genetically-defined neuronal cell types.

Neuronal shape is a fundamental property that subserves its function, but very little is known about the molecular bases of how neurons develop and maintain their shapes^86,87^. Such a paucity of knowledge is largely due to the limited scales and high cost (e.g., labor, and specialized skills or equipment) with which one can study the morphology of genetically-defined neurons in the mammalian brain. As a proof-of-concept on the utility of dendritome mapping to study the biology and disease of genetically-defined single neurons, our study uncovered aging- and HD-associated changes in D1- and D2-MSN dendrites. We showed aging-induced MSN atrophy is comparable between D1- and D2-MSNs; while for HD, separating D1- and D2-MSNs can help uncover length and complexity related pathologies, especially for D1-MSNs at the CPr level. Since the previous study of HD MSN dendritic pathology with only dozens of MSNs in the CPi region revealed inconsistent findings, one with^32^ and the other without^88^ significant dendritic changes in HD. Thus, our study clearly demonstrates the value of studying large number of completely reconstructed genetically defined neurons throughout the brain and with multiple replicates.

In summary, the dendritome mapping approach provides a novel end-to-end systems biology solution using brainwide single-neuron dendritic morphologies as readouts to uncover the genetic and environmental factors that influence the development, plasticity, and pathology of single neurons. Since all the reconstructed neurons from different studies will be analyzed in the same 3D CCF boxes, our approach will help the field to build a cumulative knowledge base for single neuron morphologies and to integrate those morphologies with other data types mapped to the same CCF space (e.g., transcriptome, axonal connectivity). Looking into the future, the dendritome mapping approach can constitute an important new systems biology approach to investigate the single-neuron biology and pathology, and for testing novel therapeutics to correct such pathology in the mammalian brain.

## ONLINE METHODS

### iDISCO+ tissue processing and imaging of thick-sectioned MORF3 tissue

Tissue dissected from MORF3 mice that were transcardially perfused with 0.1M phosphate buffered solution (PBS) and fixed with 4% paraformaldehyde (PFA) were vibratome sectioned (Compresstome, Precisionary Instruments) at 500µm. The thick-sectioned tissue was cleared and immunolabeled using our published protocol for iDISCO+ processing adapted for MORF3 tissue^8,42^. Reagents for iDISCO+ immunolabeling were updated to specifically optimize imaging of Camk2a-CreER/MORF3/D1-tdTom tissue: Sections were incubated in 1) primary antibody solution of chicken polyclonal anti-V5 tag (Bethyl; 1:500) and rabbit polyclonal anti-RFP (Rockland; 1:500), 2) fluorescent-conjugated secondary antibody solution of goat anti-chicken, Alexa Fluor Plus 488 (Invitrogen; 1:500) and goat anti-rabbit, SeTau-647 (Jackson ImmunoResearch; SETA BioMedicals; conjugated in-house according to manufacturer’s protocol), and 3) a fluorescent Nissl stain (NeuroTrace Red, Invitrogen; 1:300). iDISCO+ cleared and immunolabeled tissue were index-matched and mounted in dichloromethane (DBE) on glass microscope slides with custom-made silicone spacers (Grace Bio-Labs) to accommodate the 500µm-thick tissue for coverslipping and sealing silicone.

Sections were imaged using an Andor DragonFly spinning disk confocal (Oxford Instruments) equipped with low-power 10x and high-power 30x (0.8mm working distance) silicone immersion objectives (Olympus). For both low and high-power imaging, individual image tiles (Imaris format, .ims) were stitched into a larger montage with Imaris Stitcher (Oxford Instruments) in the Imaris format for the downstream image reconstruction pipeline.

### Computational pipeline to reconstruct neuronal morphology from 3D image volumes from thick-sectioned MORF tissue

#### Semantic segmentation of neuron soma and neurites

We trained a Convolutional Neural Network (CNN)-based model to label each voxel as one of three classes: BACKGROUND, SOMA and NEURITES. The model architecture is modified from U-Net^89^, and consists of a downsampling path, an upsampling path, and skip connections in between. The 3D model receives a sequence of image patches from adjacent z planes, and makes dense classification for the center patch.

We typically use image patch sequence with [sequence_length, height, width] equal to [3, 256, 256], which is much smaller than the whole image volume. To segment the entire image volume, we slide the 3x256x256 input window along the x, y and z axes of the volume, and tile the segmentation results together. Within each input window, we only take the center crop (128x128) of the segmentation results, where the model predictions are unaffected by image content cut-offs near the window boundary. Each input window overlaps with its neighbor window to provide full segmentation coverage for the volume.

We developed a workflow of data annotation for model training. First, some candidate image regions with high average intensity (indicating presence of fluorescent neurons) are selected from representative image volumes. Each image is annotated by an annotator, marking all SOMA and NEURITES class voxels. A second annotator then verifies the annotation for quality control.

#### Neuron cluster tracing

We use the APP2 algorithm^48^ to trace the segmented image volume. APP2 assumes a single neuron is present in the image volume, and traces this neuron starting from the center of the presumed soma. Since our image volume contains numerous neurons, the assumption of the APP2 algorithm poses two issues: (1) the algorithm does not fully trace all neurons (2) the algorithm fuses together neurons with proximate neurites. To address the former, we compute the centroids of segmented somas via connected component analysis, and run the APP2 algorithm multiple times over the volume, each time starting from a soma not covered by traces from previous runs of the algorithm. This allows us to exhaustively trace all neurons in the image volume, where some traces are neuron clusters (fusions of multiple neurons) rather than individual neurons.

#### Segment individual neurons from clusters

We use the G-Cut algorithm^47^ to segment individual neuron morphology from APP2 traced neuron clusters. APP2 presents traced morphology as directed acyclic graphs (DAG), where the nodes are sampled locations within the soma or along the neurites. The G-Cut algorithm takes as input the APP2 produced DAG, as well as centroid coordinates of segmented somas in the neuron cluster. The algorithm then assigns each neurite to one of the somas, optimizing the morphological likelihood over all possible assignments. Following the G-Cut algorithm, we obtain individual neuron morphology reconstructions for the entire image volume.

#### Single neuron cropping to facilitate manual correction and validation

We verified the pipeline neuron reconstructions manually using neuTube^49^, a semi-automatic tracing software with strong support for image volume visualization and SWC trace editing. NeuTube loads a maximum of 2GB of raw image and as such, requires cropping the image volume to single neurons for manual correction and validation using NeuTube. To automate this single-neuron cropping, we created a python script called Benchmark Manager. For each SWC trace, we programmatically crop out a local image volume that contains this single neuron, and apply necessary node coordinate offset to align the SWC trace and the local image volume. Human annotators can then examine the pipeline reconstruction and make any corrections if needed in NeuTube, and export the quality controlled and validated final reconstruction.

### CCF registration of thick-sectioned tissue

#### Resampling

We performed image registration at approximately 50-micron resolution. Downsampling was achieved by identifying an integer multiple of the pixel size that would give desired resolution, breaking it down into its prime factors, and downsampling from the smallest to largest factor using local averaging. Moving from smallest to largest factors in this manner reduces computation time.

#### Atlas to 10x sequence registration

We modeled the voxel values in our dataset as a sequence of unknown transformations applied to an atlas image. The parameters in these transformations were iteratively estimated using maximum a posteriori estimation, with missing data (artifacts and missing tissues) modeled in an Expectation Maximization setting^90,91^.

1. The atlas image is first deformed using a diffeomorphism *φ* constructed by integrating a smooth velocity field in the LDDMM setting^92^. This represents the overall shape difference between the atlas and the brain being imaged.
2. A 12 degree of freedom affine transform, *A*, is applied. This represents macroscopic changes due to tissue processing, as well as the orientation of the tissue relative to its sectioning plane.
3. A fixed (not estimated) translation, *T_j_*, is applied parallel to the z axis for each block. This represents a translation applied to the tissue block as consecutive slabs are sliced off.
4. A contrast transformation, *p*, is applied to the atlas image, zeroing out imaging data outside of the 500µm slab. This transformation of imaging data, rather than shape, as part of our model has been described as projective LDDMM^93^.
5. A second diffeomorphism, *ϕ_j_*, is applied independently to each slab. This represents shape change due to cutting or transporting the sample, in a manner that is independent from one slice to another.
6. A second 12 parameter affine, *B_j_*, transformation is applied independently to each slab. This represents macroscopic changes due to processing, as well as orientation of the tissue relative to the microscope.
7. A contrast transformation, *f_j_*, is applied to each voxel in the transformed atlas image. This represents the fact that tissue contrast being imaged is not the same as that in the atlas. These unknown contrast changes were described in^91^.

For an atlas image *I*, and a sequence of target sections *J_j_*, the objective function we minimize is:

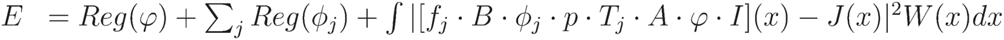

Here the · symbol is used to mean “acts on”’, and may either transform the image’s shape, or its pixel intensities. Our algorithm is implemented in pytorch for GPU parallelization and automatic gradient calculation. Velocity field parameters are estimated by Hilbert gradient descent, with regularization functions *Reg* as described in^92^. Affine transforms are estimated by Riemannian gradient descent, with the Riemannian metric described in^94^. In addition to accelerating optimization, this strategy makes our optimization invariant to units (i.e. microns versus millimeters) and placement of the origin. Weights *W* are estimated to avoid overfitting missing tissue or artifacts. They are equal to the posterior probability that each voxel represents “good” (i.e. not an artifact or missing tissue) data, and is updated using Gaussian mixture modeling as described in^91^. Note that for any hemisphere brains, we zeroed out the appropriate half of our atlas images before registration.

Due to the fact that this is a nonconvex optimization problem and we use gradient-based optimization, the alignment results exhibit a sensitivity to initialization. We found this effect to be particularly strong along the anterior-posterior axis. We developed an initialization procedure to choose an initial affine transformation that would minimize the sum of square error between the center of mass of each caudoputamen structure in the atlas, and the caudoputamen anatomical structures our experimental team believed to be in each slab. We use the Nelder-Mead procedure in the scipy package for this optimization.

#### 10x to 30x registration

Each section imaged at 10x is aligned to one or two smaller fields of views imaged at 30x. For this task we use an existing diffeomorphic registration pipeline with missing data and contrast differences, described in^45,59,91^. This algorithm computes another diffeomorphism and 12 parameter affine transform to align the two datasets. We note that due to the limited fields of view in the 30x images, older algorithms such as^92^, or symmetric algorithms such as^62^, would not be appropriate.

#### Mapping neurons to anatomical labels

Soma locations largely confined to the CP were annotated in each 30x image. These were mapped by composing with the inverses of the transformations computed above. For this reason “inverse consistency” of the transformations we estimate is critical, which motivates our choice of the LDDMM model. To evaluate the accuracy of our pipeline, each mapped neuron was assigned a probability distribution of possible regions in the in the Unified atlas (anatomical labels from both the Franklin-Paxinos (FP) and the common coordinate framework (CCF) from the Allen Institute for Brain Science)^59^. Many of the anatomical regions were obtained by drawing straight cutting planes or clustering voxels into neighborhoods, rather than relying on contrast differences that can be seen in imaging data, or actual projection studies. For this reason, when neurons are mapped close to a boundary we have a low confidence about which region they actually belong to, and we use a “soft assignment’” approach. A distance transform was applied to binary masks for each anatomical region of interest, and exponentially decaying tails with length scale 5μm were computed. These were interpreted as probability densities over space that represented the prior probability of a neuron’s location, given knowledge of only its structure id. This was combined with the actual position of our mapped neuron using Bayes theorem to produce a posterior distribution. We used the regions with the top 3 posterior probabilities to compare to an anatomist’s region assignment on a subset of slices. Our agreement is illustrated in Fig. 2d.

#### Unbiased analysis of neuron distribution by mapping to a coarse grid

Since the anatomical labels may not be “true”, we complement our above analysis with an unbiased approach to quantify cell density as a function of space, using their (x,y,z) locations (with left side neurons reflected to match right) rather than their region assignments. We divide the Allen CCF into 500µm cubic boxes, and give each a unique integer id, with left side and right side corresponding boxes differing only by a sign, leading to 7020 boxes. We focus our analysis on boxes that contain some portion of the CP, which included 465 boxes. An example section is shown in Fig. 5a

#### Interactive visualization with Neuroglancer

Using Neuroglancer, a web-based visualization framework of volumetric data, we display several mouse brain atlases along with the registration results for one registered brain. In this view, we overlay the Allen mouse brain atlas, Yongsoo Kim’s anatomical labels, one registered mouse brain from the P56 dataset, and the soma locations of the registered neurons associated with this brain as illustrated in Fig. 2c.

### Post-processing of reconstructions

#### Post processing of reconstruction and morphometric extraction

The neuronal reconstructions obtained from the in-house reconstruction pipeline are often incomplete and need further processing. Therefore, the automatically generated 3D tracings were processed for its completeness and necessary corrections by a well-trained reconstruction team using neuTube^49^. The proofread reconstructions were then visually inspected and validated by expert scientists to ensure the accuracy of structure and branching patterns. During the manual reconstruction process, some errors are overlooked due to the limitations of reconstruction software tools, therefore, a final quality control check was implemented using a series of semi-automated programs developed in-house and also provided by the NeuroMorpho.Org team. These tools further improved the 3D neuronal reconstruction quality by eliminating any spurious long branches, soma and dendrites tag inconsistencies (i.e. actual tags of soma =1, basal dendrites =3, respectively), trifurcations, overlapping branches, small side branches, and node radii irregularities. Since the benchmark manager of the reconstruction pipeline crops each brain section image into smaller crops based on cell body location and individual neurons, the post-processed reconstructions were then mapped to the original 30x image space and properly scaled. Finally, morphometric features were extracted and calculated using L-measure V5.3^50,51^ and the TREES Toolbox^52^. Out of 35 L-Measure morphometric parameters, 7 features including diameter, surface, volume, helix, contraction, fragmentation, and pk classic were excluded because these features are more prone to errors and variations due to their dependence on tissue processing, imaging modalities, varying signal intensities, and tracing software tools. The remaining 31 morphometric parameters describe length-/size-related, complexity-related, and angle-related characteristics, and were used to perform the statistical analyses.

#### D1- and D2- type assignment

The classification of D1 and D2 medium spiny neurons (MSNs) was conducted using a machine learning approach based on soma signal intensity in D1-TdTomato channel images. A dataset of 300 D1-TdTomato channel images was randomly selected, with 80% allocated for training and 20% for testing. Images were cropped to focus on the soma region, and a Gaussian filter was applied for preprocessing. Three classifiers, Support Vector Machine (SVM)^66^, Random Forest^95^, and Convolutional Neural Network (CNN)^96^, were trained separately. For the SVM and Random Forest models, hyperparameter tuning was performed using grid search with 3-fold cross-validation. The CNN model was optimized using data augmentation techniques and early stopping to prevent overfitting. The model assignment of D1- and D2-MSNs was determined by ensemble averaging the prediction probabilities from the three models. The model predictions were subsequently referenced by human experts to determine the final assignment of D1 and D2 MSNs.

### Statistical analysis

#### Morphological classification of P56 D1/D2- MSNs and Morphological Territories (MT)

Principal component analysis of P56 D1/D2- MSNs and Morphological territories was performed to reduce the redundancy among morphological parameters^97^. First 15 principal components which contributed more than 95% of the variance were then utilized to perform supervised machine learning classification. We leveraged support vector machine^66^, a binary classification algorithm that gives the helps separating data points of different classes in multidimensional space by finding the maximum margin hyperplane, with 10-fold cross validation repeated 5 times for the categorization of P56 D1- vs D2- MSNs, D1- vs D2- MSNs in striatal subregions (CPr CPi and CPc), as well as the categorization three main striatal subregions with all D1- and D2- MSNs combined. Similarly, the binary classification of the morphological territories (e.g., MT-1 vs MT- 2, MT-1 vs MT-3, etc.) was also performed using SVM and 10-fold cross validation with 5 repeats. This analysis was performed in R 4.2.2.

#### Morphological comparisons of P56 and 12-m MSNs

The grouping of morphometric features was first studied based on their biweight midcorrelation and the hierarchical clustering tree was cut at the height of 0.15 to group the highly-correlated features together. The comparison of morphometric differences of D1- and D2- MSNs in all regions combined and within each striatal subregions/communities were conducted using two sample t-test. The regional variations in MSN morphology across the 3 striatal subregions were also analyzed using the two sample t-test, with morphometric features normalized to the corresponding median values within each brain to facilitate the comparison across striatal subregions and between brains on the same scale. The differences of Huntington’s disease (HD) compared to WT in 12-m of age and aging effect (12-m WT vs. P56 WT) were assessed using two sample t-test, and the effects were tested in all MSNs, D1- MSNs and D2- MSNs in 3 striatal subregions (CPr, CPi and CPc). All p-values were adjusted for multiple comparisons using the Bonferroni method. The analyses were performed in R 4.2.2.

#### Unbiased box-based analysis of P56 MSNs

In box-based analyses, 21 non-redundant features were selected based on the feature clustering result. The features were then centered and scaled, the scaling (to a mean of 0 and variance of 1) ensures that differences in scale and variation of the morphometric measures do not bias the principal component toward specific measures. Then, the “eigen-morph” for each box was calculated using the scaled matrix of morphometric features of the neurons in the boxes and the *ModuleEigengenes()* function from Weighted Gene Co-expression Network Analysis (*WGCNA*)^38^ was used to create a synthetic descriptor of each box, which is basically the first principal component of morphometrics of the group of neurons in the box that summarizes the collective behavior of neurons within each box. The “eigen-morph” can be viewed as a summary or prototype shape of the neurons in each box. Subsequently, a box-box pairwise dissimilarity measure was calculated to form a dissimilarity matrix and then used to cluster the boxes using hierarchical clustering. Various arguments of *DynamicTreeCut()* were applied to cut the dendrogram at different levels, with the final clusters determined by considering the Silhouette score and the best biological interpretations. After the clusters were formed, the connected components and their sizes were identified using the distances of centroids between each box, and the distribution of the sizes of connected components was tested against the uniform distribution (chi-square test). The morphological territories were defined as the connected components within each cluster that has more than 7 connected boxes. The representative neurons were selected based on the contributions to the first principal component of each morphological territory. The analyses were performed in R 4.2.2.

## ACKNOWLEDGEMENTS

This research was supported by the National Institutes of Health’s (NIH’s) BRAIN Initiative grants to X.W.Y. and H.W.D (U01MH117079) and X.W.Y., H.W.D., and D.T. (RF1MH128888). The HD and aging related work in this study were funded by CHDI Foundation, Inc. X.W.Y. is also supported by NINDS/NIH grants (R01NS113612), the Terry Semel Chair in Alzheimer’s Disease Research and Treatment at UCLA, the Hereditary Disease Foundation. Yongsoo Kim is funded by NIH RF1MH124605. We acknowledge the pivotal contribution of ver 40 undergraduate students from UCLA and other universities as well as high school students who performed the manual neuronal reconstruction and proofreading for the Yang Lab. We are indebted to Zhuhao Wu (The Weill Cornell Medical College) for the iDISCO+ protocol.

## AUTHOR CONTRIBUTIONS

X.W.Y. conceptualized and directed the study. X.W.Y. and C.S.P. designed the experiments and C.S.P. managed the entire project. X.W.Y., C.S.P., and M.Y., interpreted the data and wrote the manuscript. M.A.A., P.L., H.W.D., N.N.F, D.T., and A.B. contributed to the interpretation and writing of specific sections and creation of related figure panels. D.T. and A.B. developed the multiscale tissue section CCFv3 registration pipeline and created Neuroglancer visualization. C.S.P. and H.W.D. developed the imaging pipeline. J.Y.Z. and R.V. provided technical support. M.Z., H.W.D., C.S.P. developed the reconstruction pipeline with support from K. Marrett, J.C., and K. Moradi. Imaging processing was done by C.S.P., J.Y.Z., C.C., M.A.A., and G.M. Neuronal reconstruction and validation were performed by C.C., M.A.A. G.M. and K.W. and the Yang Lab Neuronal Reconstruction Team (40 mostly UCLA undergraduate students). The latter included S.N., K.W., J.R.Z., S.L., C.S., M.J.L., M.L., who had produced the most reconstructed neurons in our dataset. Statistical analysis of neuronal morphometrics and related figure plots were made by M.Y., with contribution from P.L., M.A.A. and S.N. G.A.A. provide consultation on morphometric analyses. Y.K. provided the fine striatal anatomical map in CCFv3. X.W.Y. conceptualized and M.Y., D.T., and A.B. developed the Box and Eigen-morph analyses.

## COMPETING INTERESTS

X.W.Y. has served on Scientific Advisory Board for Lyterian Therapeutics, Ophidion Therapeutics, Function Rx, Triplet Therapeutics, and Mitokinin, Inc.; and has consulted for Ionis, Biogen, Novartis, Roche, PTC Therapeutics, Sangamo, LifeEdit, Ascidian, Forbion, and ReviR. P.L. serves as occasional bioinformatics consultant for The Bioinformatics CRO, Inc; Vynance Technologies, LLC; and FOXO Technologies, Inc.

**Supplementary Table 1. Description of morphometric features.**

**Supplementary Table 2. Raw morphometrics for all reconstructions from P56 WT, 12m WT, and 12m Q140 brains.**

**Supplementary Table 3. Abbreviations of corticostriatal projection domains in dorsal striatum.**

**Supplementary Table 4. Individual registered neuron level/community assignment probability for WT P56, 12m WT, and 12m Q140 brains.**

**Supplementary Table 5. Support vector classification results for P56 WT subtype and regional (D1- vs D2-MSN and rostral-caudal level) comparisons and comparisons between box-analysis morphometric territories (MTs).**

---

**Supplementary Fig. 1.**
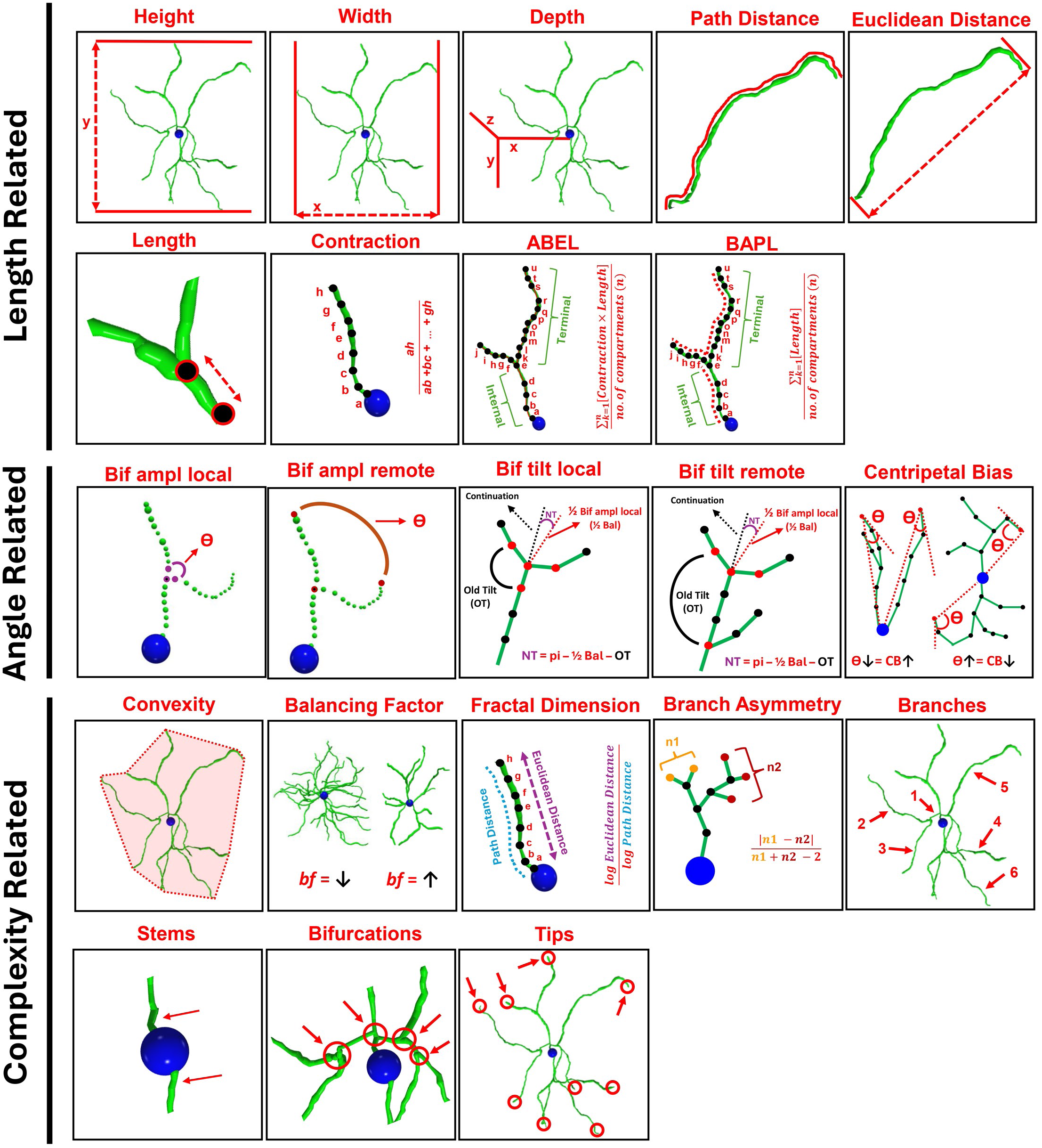
Illustrated summary of morphometrics. Representation of the main morphometric features used in this study grouped into 3 main structural characteristics including length related (height, length, etc.), angle related (bifurcation amplitude local, centripetal bias, etc.), and complexity related (convexity, bifurcations, etc.). Detailed explanation and abbreviations of all the morphometric parameters can be found in Supplementary Table 1.

**Supplementary Fig. 2.**
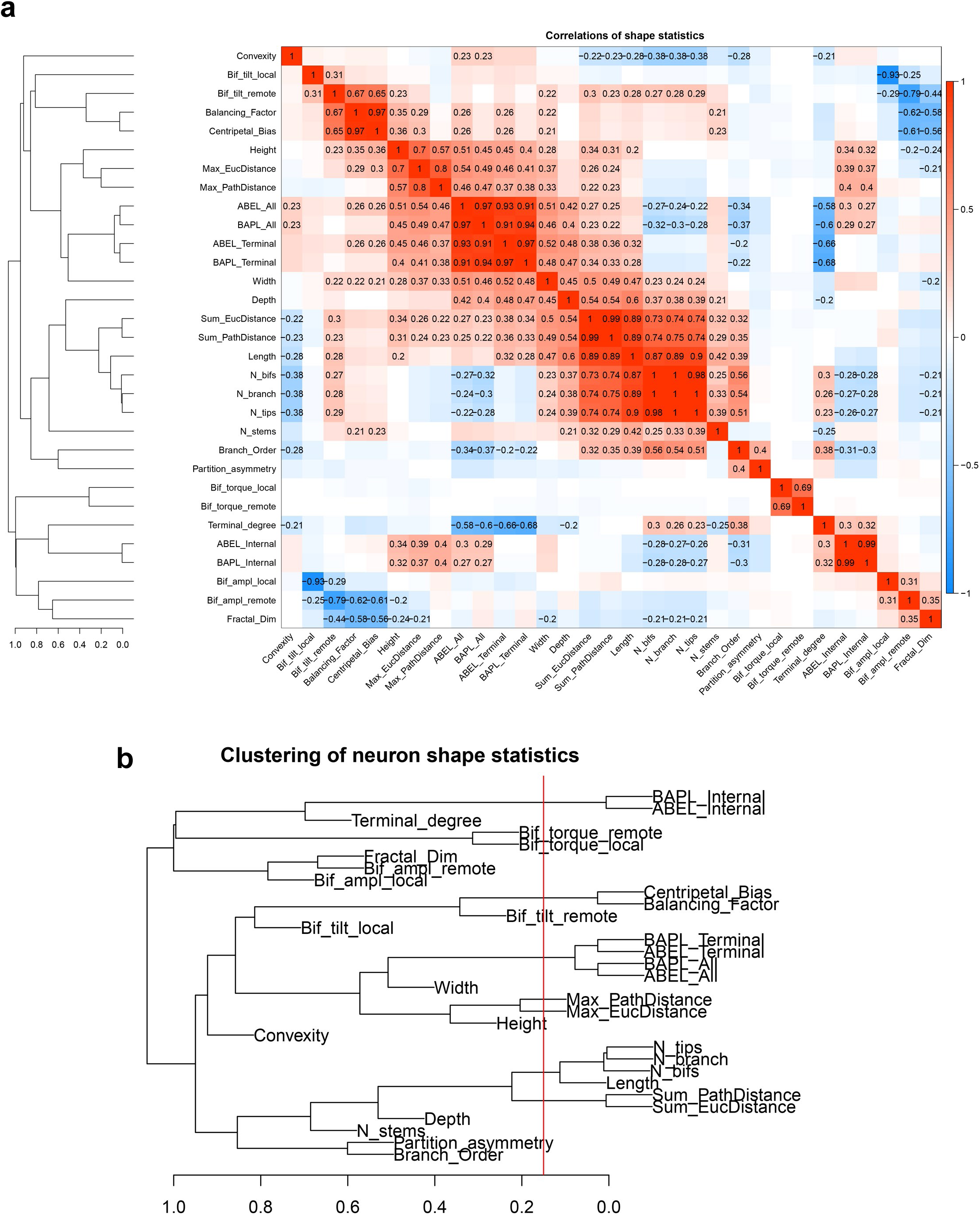
Correlation of morphometric features and clustering. **a,** Heatmap of bi-weight mid-correlation between 31 morphometric features; red denotes positive correlations and blue denotes negative correlations. **b,** The hierarchical clustering results of 31 morphometric features, with tree cutting at height = 0.15 to group the highly-correlated morphometrics.

**Supplementary Fig. 3.**
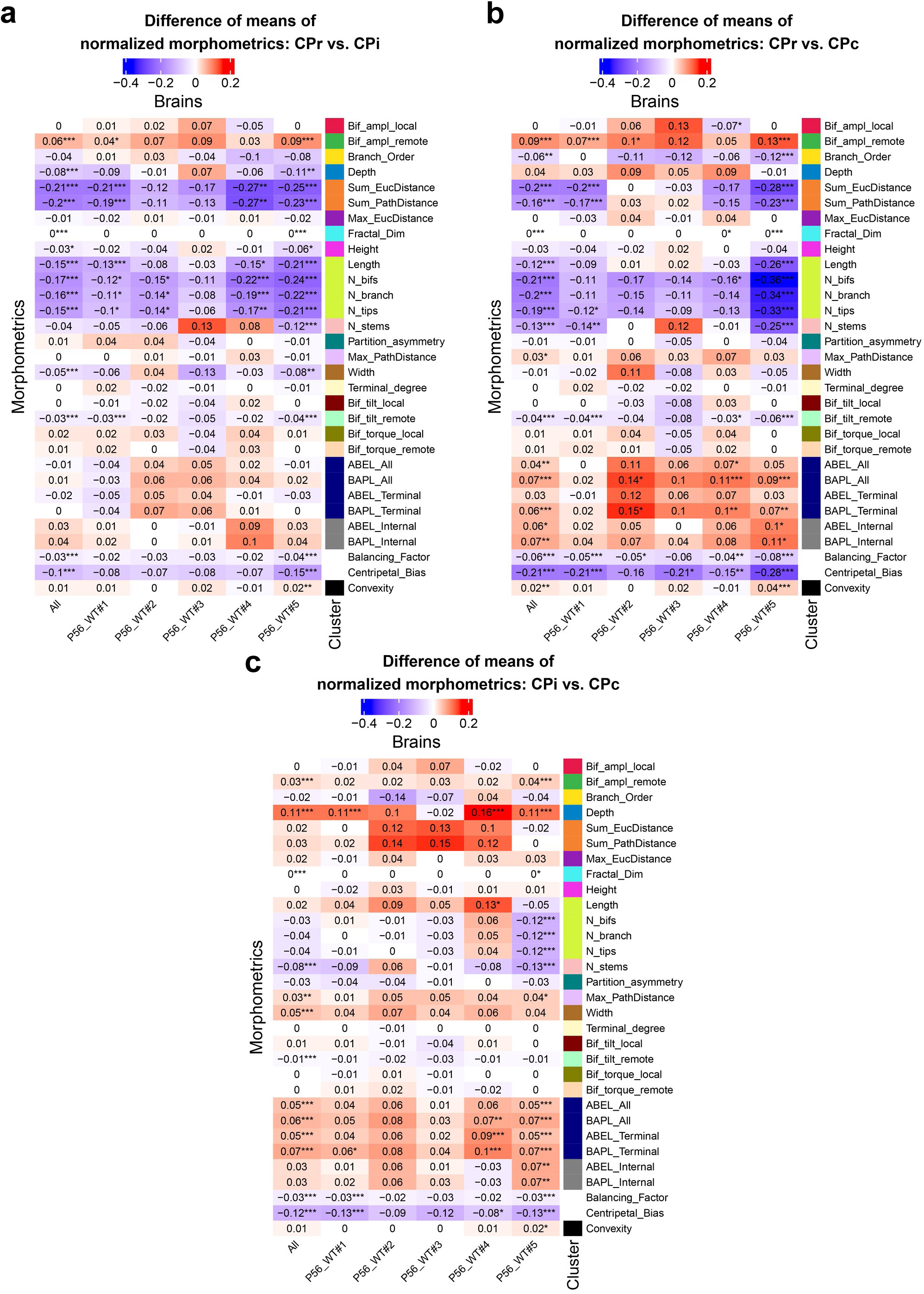
Heatmap distribution of normalized morphometrics for all MSNs in P56 WT brains. 3 heatmaps for subregion pairwise comparisons (**a,** CPr vs. Cpi; **b,** CPr vs. CPc; and **c,** CPi vs. CPc) for all P56 WT brains (n=5) combined and by individual brains (two- sample t-test was performed). Blue denotes values are lower than the brain median and red denotes values are higher than the brain median.

**Supplementary Fig. 4.**
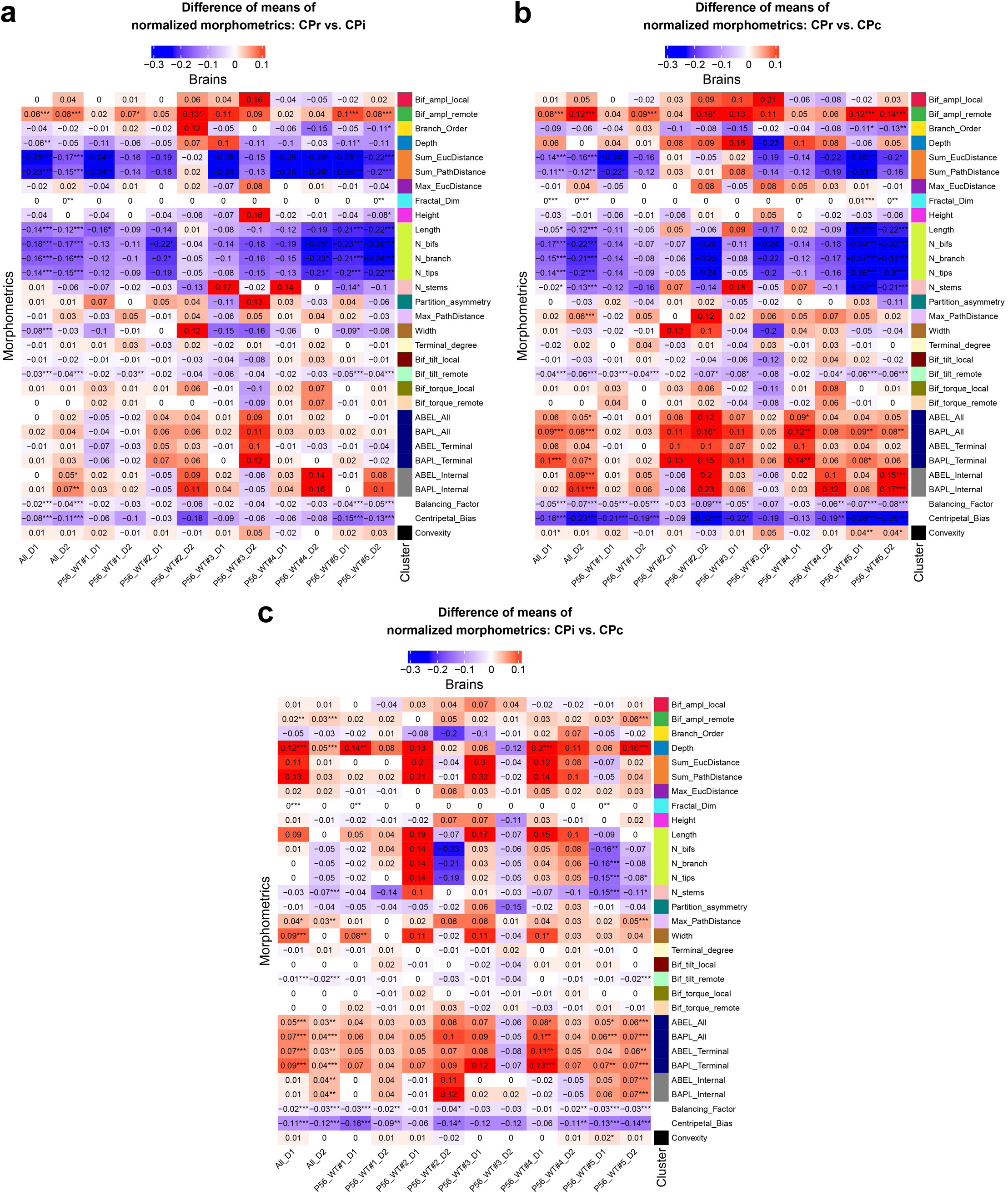
Heatmap distribution of normalized morphometrics for D1- and D2- MSNs in P56 WT brains. 3 heatmaps for subregion pairwise comparisons (**a,** CPr vs. Cpi; **b,** CPr vs. CPc; and **c,** CPi vs. CPc) in all P56 WT brains (n=5) combined and by individual brains, separated by D1- and D2- MSNs (two-sample t-test was performed). Blue denotes values are lower than the brain median and red denotes values are higher than the brain median.

**Supplementary Fig. 5.**
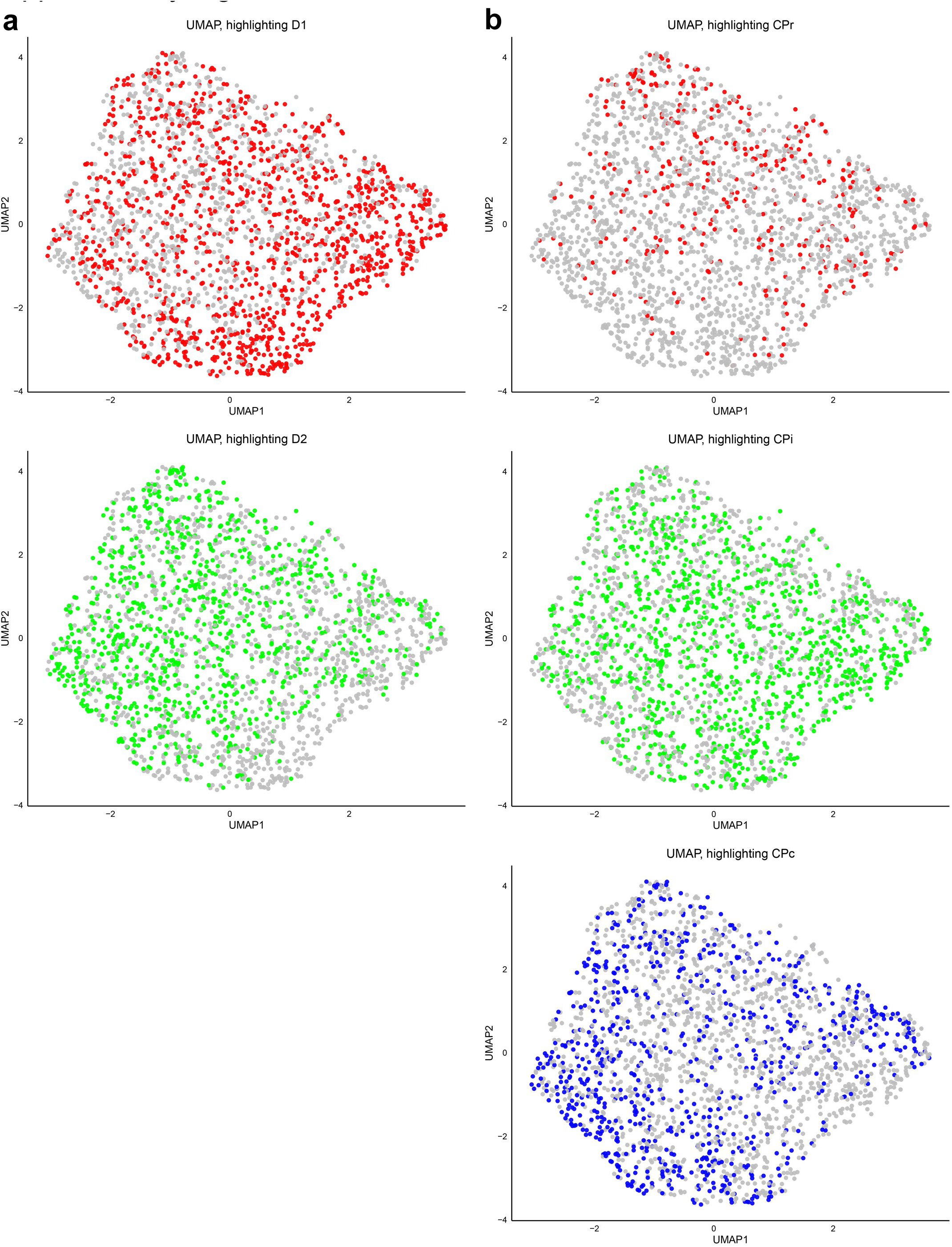
UMAP (UMAP1 vs. UMAP2) plots of D1, D2, CPr, CPi, CPc. **a,** The UMAP showing the distribution of D1- MSNs (red) and D2- MSNs (green) separately. **b,** The UMAP showing the distribution of CPr cells (red), CPi cells (green) and CPc cells (blue) separately.

**Supplementary Fig. 6.**
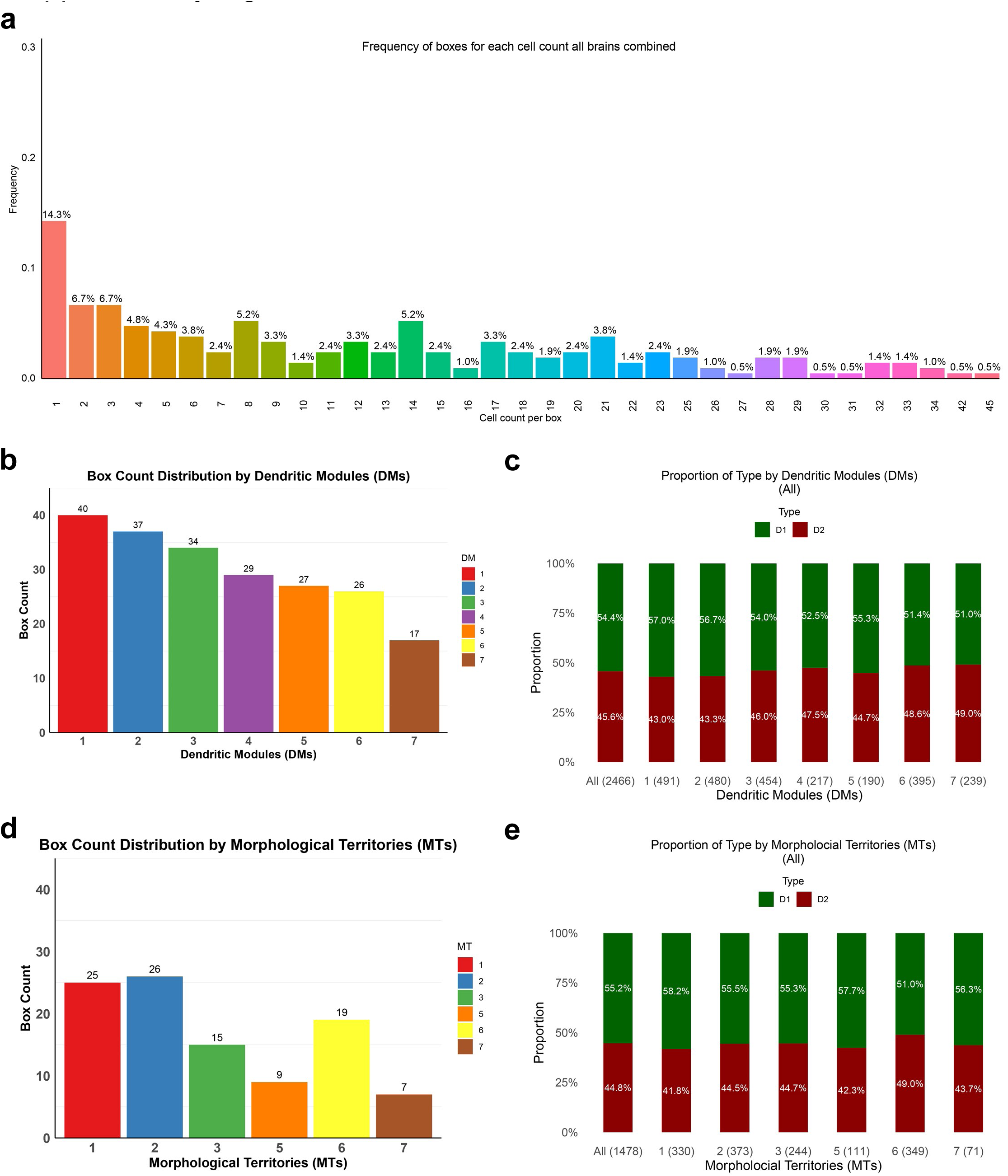
Summary statistics for box analysis dendritic modules (DMs) and morphometric territories (MTs). **a,** The frequency distribution of box cell counts, box cell counts ranges from the least of 1 cell per box to the max of 45 cells per box. **b,** The distribution of box counts in each DM after the clustering of the boxes. **c,** The D1- and D2- MSN proportion distribution, overall and within each DM, the proportions are relatively even (chi-square goodness-of-fit test was performed). **d,** The distribution of box counts in MT after filtering for connected boxes (>7) in each DM. **e,** The D1- and D2- MSN proportion distribution, overall and within each MT, the proportions are relatively even (chi-square goodness-of-fit test was performed).

**Supplementary Fig. 7.**
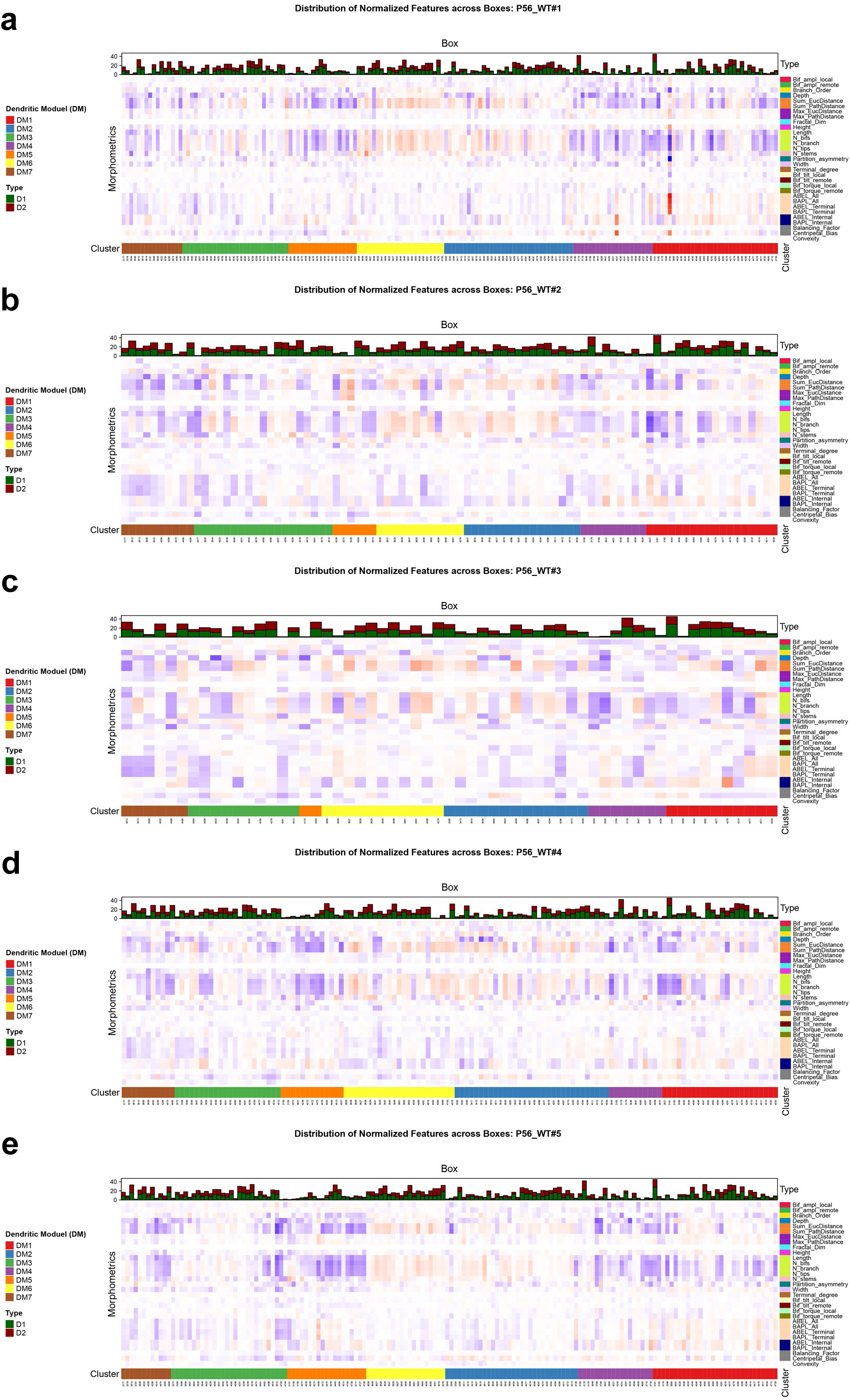
Distribution of normalized features across boxes within each dendritic module (DM). 5 heatmaps for each P56 WT brain (**a-e**) in which boxes were grouped and ordered by clustering results, with D1- and D2- MSNs distribution per box; red denotes value above the median and blue denotes value above the median.

**Supplementary Fig. 8.**
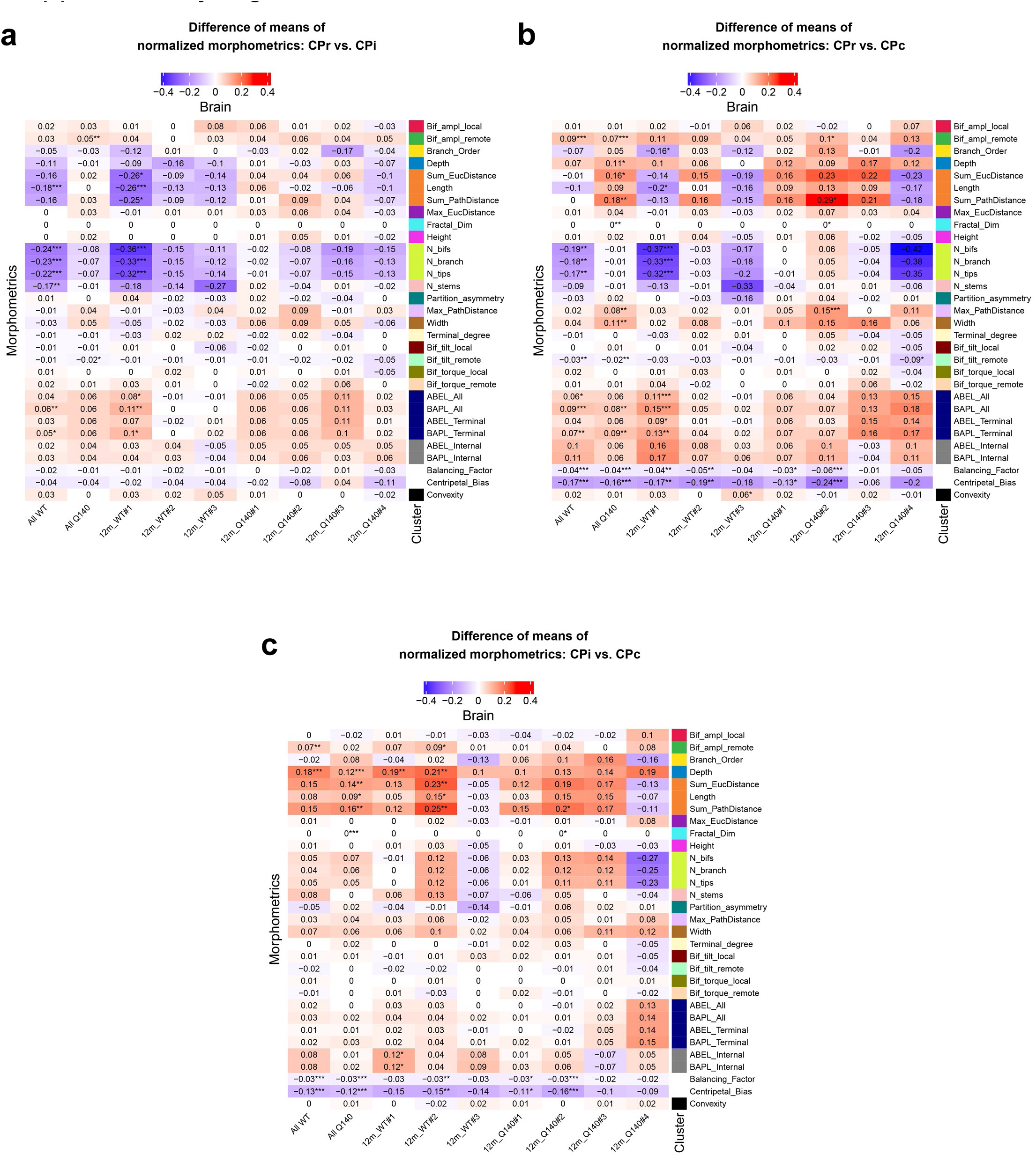
Heatmap distribution of normalized morphometrics for all MSNs in 12m WT and Q140 brains. 3 heatmaps for subregion pairwise comparisons (**a,** CPr vs. Cpi; **b,** CPr vs. CPc; and **c,** CPi vs. CPc) for all 12m WT (n=3) and 12m Q140 (n=4) brains combined by genotype and by individual brains (two-sample t-test was performed). Blue denotes values are lower than the brain median and red denotes values are higher than the brain median.

**Supplementary Fig. 9.**
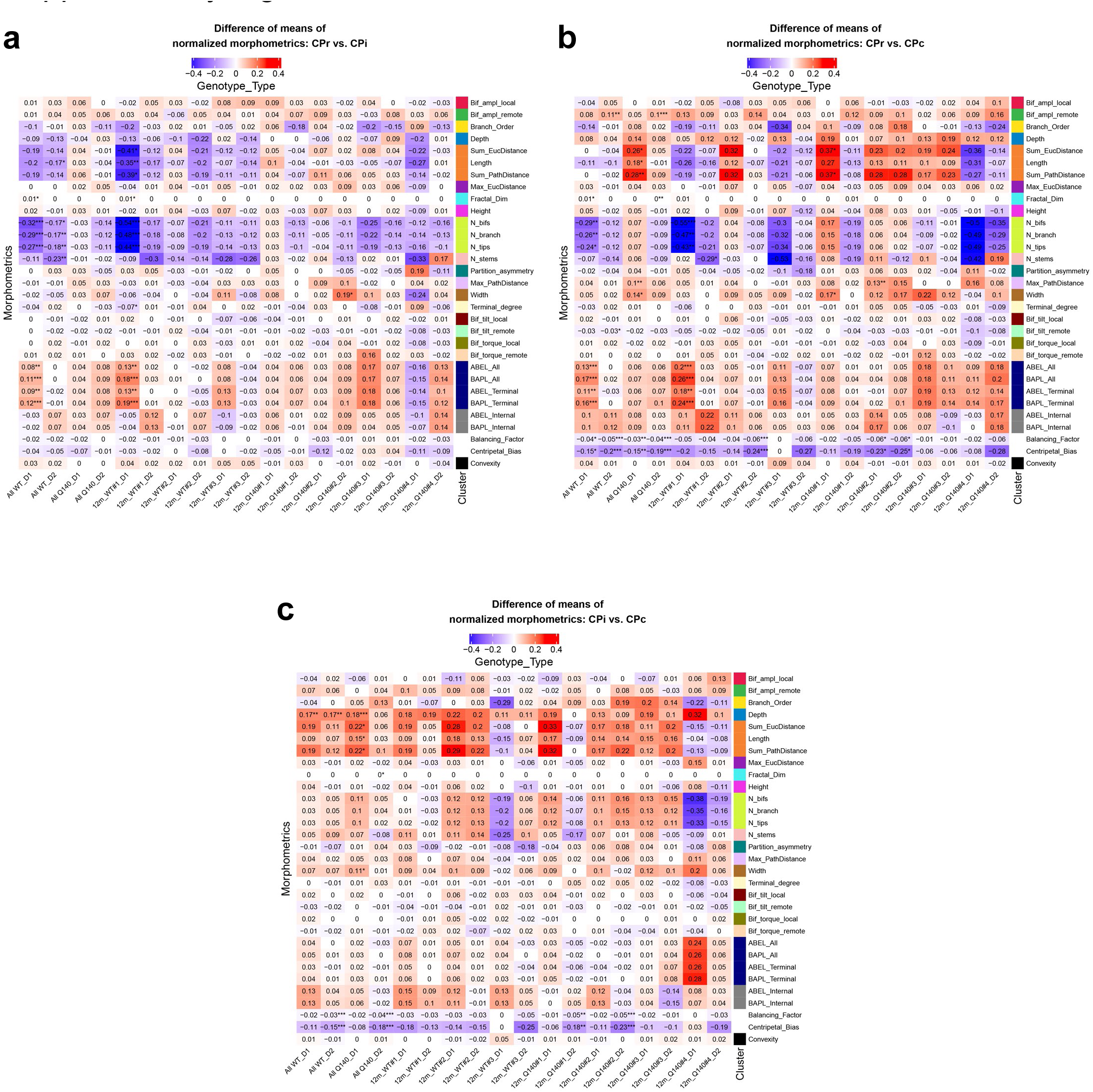
Heatmap distribution of normalized morphometrics for D1- and D2- MSNs in 12m WT and Q140 brains. 3 heatmaps for subregion pairwise comparisons (**a,** CPr vs. Cpi; **b,** CPr vs. CPc; and **c,** CPi vs. CPc) for all 12m WT (n=3) and 12m Q140 (n=4) brains combined by genotype and by individual brains, separated by D1- and D2- MSNs (two-sample t- test was performed). Blue denotes values are lower than the brain median and red denotes values are higher than the brain median.

**Supplementary Fig. 10.**
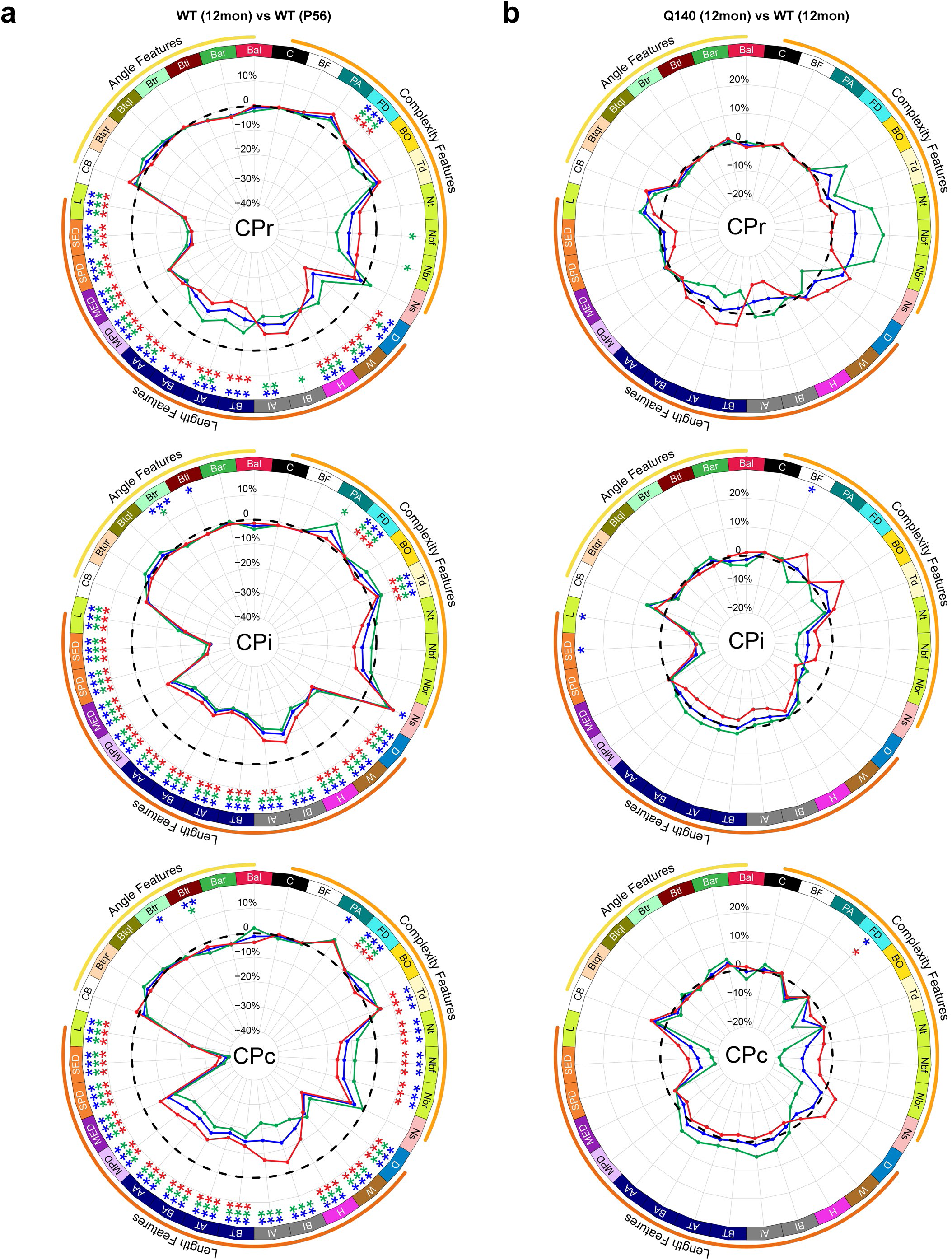
Radar plots for 12m WT vs P56 and 12m WT vs 12m Q140 at CPr, CPi, and CPc striatal levels. **a**, The percentage changes of morphometric features of MSNs from 12m WT (n=3) compared to P56 WT mice (n=5) across the 3 rostral-caudal levels (two- sample t-test was performed). **b**, The percentage changes of morphometric features of MSNs from 12m Q140 (n=4) compared to 12m WT mice (n=3) across the 3 rostral-caudal levels (two- sample t-test was performed). *P < 0.05, **P < 0.01, ***P < 0.001; in **a** and **b**, blue asterisks for both D1- and D2-MSNs, green asterisks for D1-MSNs only, and red asterisks for D2-MSNs only. Morphometrics abbreviations: 1) Bal, Bif_ampl_local; 2) Bar, Bif_ampl_remote; 3) Btl, Bif_tilt_local; 4) Btr, Bif_tilt_remote; 5) Btql, Bif_torque_local; 6) Btqr, Bif_torque_remote; 7) CB, Centripetal_Bias; 8) L, Length; 9) SED, Sum_EucDistance; 10) SPD, Sum_PathDistance; 11) MED, Max_EucDistance; 12) MPD, Max_PathDistance; 13) AA, ABEL_All; 14) BA, BAPL_All; 15) AT, ABEL_Terminal; 16) BT, BAPL_Terminal; 17) AI, ABEL_Internal; 18) BI, BAPL_Internal; 19) H, Height; 20) W, Width; 21) D, Depth; 22) Ns, N_stems; 23) Nbr, N_branch; 24) Nbf, N_bifs; 25) Nt, N_tips; 26) Td, Terminal_degree; 27) BO, Branch_Order; 28) FD, Fractal_Dim; 29) PA, Partition_asymmetry; 30) BF, Balancing_Factor; 31) C, Convexity.

## REFERENCES

1. Ramón y Cajal, S. & Ramón y Cajal, S. Histology of the Nervous System of Man and Vertebrates. (Oxford Univ. Press, New York, 1995).

2. BRAIN Initiative Cell Census Network (BICCN) et al. A multimodal cell census and atlas of the mammalian primary motor cortex. Nature 598, 86–102 (2021).

3. Yao, Z. et al. A high-resolution transcriptomic and spatial atlas of cell types in the whole mouse brain. Nature 624, 317–332 (2023).

4. Adams, A. et al. International Brain Initiative: An Innovative Framework for Coordinated Global Brain Research Efforts. Neuron 105, 212–216 (2020).

5. Langlieb, J. et al. The cell type composition of the adult mouse brain revealed by single cell and spatial genomics. Preprint at 10.1101/2023.03.06.531307 (2023).

6. Jefferis, G. S. & Livet, J. Sparse and combinatorial neuron labelling. Curr. Opin. Neurobiol. 22, 101–110 (2012).

7. Zong, H., Espinosa, J. S., Su, H. H., Muzumdar, M. D. & Luo, L. Mosaic Analysis with Double Markers in Mice. Cell 121, 479–492 (2005).

8. Veldman, M. B. et al. Brainwide Genetic Sparse Cell Labeling to Illuminate the Morphology of Neurons and Glia with Cre-Dependent MORF Mice. Neuron 108, 111–127.e6 (2020).

9. Tecuatl, C., Ljungquist, B. & Ascoli, G. A. Accelerating the continuous community sharing of digital neuromorphology data. FASEB BioAdvances 6, 207–221 (2024).

10. Markram, H. et al. Reconstruction and Simulation of Neocortical Microcircuitry. Cell 163, 456–492 (2015).

11. Winnubst, J. et al. Reconstruction of 1,000 Projection Neurons Reveals New Cell Types and Organization of Long-Range Connectivity in the Mouse Brain. Cell 179, 268–281.e13 (2019).

12. Peng, H. et al. Morphological diversity of single neurons in molecularly defined cell types. Nature 598, 174–181 (2021).

13. Gao, L. et al. Single-neuron projectome of mouse prefrontal cortex. Nat. Neurosci. 25, 515–529 (2022).

14. Gao, L. et al. Single-neuron analysis of dendrites and axons reveals the network organization in mouse prefrontal cortex. Nat. Neurosci. 26, 1111–1126 (2023).

15. Scala, F. et al. Phenotypic variation of transcriptomic cell types in mouse motor cortex. Nature 598, 144–150 (2021).

16. Qiu, C. et al. A single-cell time-lapse of mouse prenatal development from gastrula to birth. Nature 626, 1084–1093 (2024).

17. Yin, H. *The Integrative Functions of the Basal Ganglia*. (CRC Press, Boca Raton, FL, 2024).

18. Gerfen, C. R. & Surmeier, D. J. Modulation of striatal projection systems by dopamine. Annu. Rev. Neurosci. 34, 441–466 (2011).

19. Plotkin, J. L. & Surmeier, D. J. Corticostriatal synaptic adaptations in Huntington’s disease. Curr. Opin. Neurobiol. 33, 53–62 (2015).

20. Gunaydin, L. A. & Kreitzer, A. C. Cortico-Basal Ganglia Circuit Function in Psychiatric Disease. Annu. Rev. Physiol. 78, 327–350 (2016).

21. Kreitzer, A. C. & Malenka, R. C. Striatal plasticity and basal ganglia circuit function. Neuron 60, 543–554 (2008).

22. Russo, S. J. et al. The addicted synapse: mechanisms of synaptic and structural plasticity in nucleus accumbens. Trends Neurosci. 33, 267–276 (2010).

23. Lobo, M. K., Karsten, S. L., Gray, M., Geschwind, D. H. & Yang, X. W. FACS-array profiling of striatal projection neuron subtypes in juvenile and adult mouse brains. Nat. Neurosci. 9, 443–452 (2006).

24. Heiman, M. et al. A Translational Profiling Approach for the Molecular Characterization of CNS Cell Types. Cell 135, 738–748 (2008).

25. Saunders, A. et al. Molecular Diversity and Specializations among the Cells of the Adult Mouse Brain. Cell 174, 1015–1030.e16 (2018).

26. Foster, N. N. et al. The mouse cortico-basal ganglia-thalamic network. Nature 598, 188–194 (2021).

27. Gerfen, C. R. & Bolam, J. P. The Neuroanatomical Organization of the Basal Ganglia. In Handbook of Behavioral Neuroscience vol. 24 3–32 (Elsevier, 2016).

28. Cazorla, M., Shegda, M., Ramesh, B., Harrison, N. L. & Kellendonk, C. Striatal D2 receptors regulate dendritic morphology of medium spiny neurons via Kir2 channels. J. Neurosci. Off. J. Soc. Neurosci. 32, 2398–2409 (2012).

29. Witzig, V. S., Komnig, D. & Falkenburger, B. H. Changes in Striatal Medium Spiny Neuron Morphology Resulting from Dopamine Depletion Are Reversible. Cells 9, 2441 (2020).

30. Fieblinger, T. et al. Cell type-specific plasticity of striatal projection neurons in parkinsonism and L-DOPA-induced dyskinesia. Nat. Commun. 5, 5316 (2014).

31. Dias-Ferreira, E. et al. Chronic stress causes frontostriatal reorganization and affects decision-making. Science 325, 621–625 (2009).

32. Goodliffe, J. W. et al. Differential changes to D1 and D2 medium spiny neurons in the 12- month-old Q175+/- mouse model of Huntington’s Disease. PLOS ONE 13, e0200626 (2018).

33. Klapstein, G. J. et al. Electrophysiological and morphological changes in striatal spiny neurons in R6/2 Huntington’s disease transgenic mice. J. Neurophysiol. 86, 2667–2677 (2001).

34. Graveland, G. A., Williams, R. S. & DiFiglia, M. Evidence for degenerative and regenerative changes in neostriatal spiny neurons in Huntington’s disease. Science 227, 770–773 (1985).

35. Gagnon, D. et al. Striatal Neurons Expressing D1 and D2 Receptors are Morphologically Distinct and Differently Affected by Dopamine Denervation in Mice. Sci. Rep. 7, 41432 (2017).

36. Gertler, T. S., Chan, C. S. & Surmeier, D. J. Dichotomous anatomical properties of adult striatal medium spiny neurons. J. Neurosci. Off. J. Soc. Neurosci. 28, 10814–10824 (2008).

37. Rosen, G. D. & Williams, R. W. Complex trait analysis of the mouse striatum: independent QTLs modulate volume and neuron number. BMC Neurosci. 2, 5 (2001).

38. Langfelder, P. & Horvath, S. WGCNA: an R package for weighted correlation network analysis. BMC Bioinformatics 9, 559 (2008).

39. Madisen, L. et al. A robust and high-throughput Cre reporting and characterization system for the whole mouse brain. Nat. Neurosci. 13, 133–140 (2010).

40. Shuen, J. A., Chen, M., Gloss, B. & Calakos, N. Drd1a-tdTomato BAC transgenic mice for simultaneous visualization of medium spiny neurons in the direct and indirect pathways of the basal ganglia. J. Neurosci. Off. J. Soc. Neurosci. 28, 2681–2685 (2008).

41. Muñoz-Castañeda, R. et al. Cellular anatomy of the mouse primary motor cortex. Nature 598, 159–166 (2021).

42. Renier, N. et al. iDISCO: A Simple, Rapid Method to Immunolabel Large Tissue Samples for Volume Imaging. Cell 159, 896–910 (2014).

43. Renier, N. et al. Mapping of Brain Activity by Automated Volume Analysis of Immediate Early Genes. Cell 165, 1789–1802 (2016).

44. Newmaster, K. T., Kronman, F. A., Wu, Y. & Kim, Y. Seeing the Forest and Its Trees Together: Implementing 3D Light Microscopy Pipelines for Cell Type Mapping in the Mouse Brain. Front. Neuroanat. 15, 787601 (2022).

45. Wang, Q. et al. The Allen Mouse Brain Common Coordinate Framework: A 3D Reference Atlas. Cell 181, 936–953.e20 (2020).

46. Parekh, R. & Ascoli, G. A. Quantitative Investigations of Axonal and Dendritic Arbors: Development, Structure, Function, and Pathology. The Neuroscientist 21, 241–254 (2015).

47. Li, R. et al. Precise segmentation of densely interweaving neuron clusters using G-Cut. Nat. Commun. 10, 1549 (2019).

48. Xiao, H. & Peng, H. APP2: automatic tracing of 3D neuron morphology based on hierarchical pruning of a gray-weighted image distance-tree. Bioinformatics 29, 1448–1454 (2013).

49. Feng, L., Zhao, T. & Kim, J. neuTube 1.0: A New Design for Efficient Neuron Reconstruction Software Based on the SWC Format. eneuro 2, ENEURO.0049-14.2014 (2015).

50. Scorcioni, R., Polavaram, S. & Ascoli, G. A. L-Measure: a web-accessible tool for the analysis, comparison and search of digital reconstructions of neuronal morphologies. Nat. Protoc. 3, 866–876 (2008).

51. Akram, M. A., Wei, Q. & Ascoli, G. A. Machine learning classification reveals robust morphometric biomarker of glial and neuronal arbors. J. Neurosci. Res. 101, 112–129 (2023).

52. Cuntz, H., Forstner, F., Borst, A. & Häusser, M. The TREES toolbox--probing the basis of axonal and dendritic branching. Neuroinformatics 9, 91–96 (2011).

53. Ascoli, G. A., Donohue, D. E. & Halavi, M. NeuroMorpho.Org: A Central Resource for Neuronal Morphologies. J. Neurosci. 27, 9247–9251 (2007).

54. Cannon, R. C., Turner, D. A., Pyapali, G. K. & Wheal, H. V. An on-line archive of reconstructed hippocampal neurons. J. Neurosci. Methods 84, 49–54 (1998).

55. Mehta, K. et al. Online conversion of reconstructed neural morphologies into standardized SWC format. Nat. Commun. 14, 7429 (2023).

56. Langfelder, P. et al. Integrated genomics and proteomics define huntingtin CAG length– dependent networks in mice. Nat. Neurosci. 19, 623–633 (2016).

57. Menalled, L. B., Sison, J. D., Dragatsis, I., Zeitlin, S. & Chesselet, M. Time course of early motor and neuropathological anomalies in a knock-in mouse model of Huntington’s disease with 140 CAG repeats. J. Comp. Neurol. 465, 11–26 (2003).

58. Halavi, M. et al. NeuroMorpho.Org Implementation of Digital Neuroscience: Dense Coverage and Integration with the NIF. Neuroinformatics 6, 241 (2008).

59. Chon, U., Vanselow, D. J., Cheng, K. C. & Kim, Y. Enhanced and unified anatomical labeling for a common mouse brain atlas. Nat. Commun. 10, 5067 (2019).

60. Park, J. et al. Integrated platform for multiscale molecular imaging and phenotyping of the human brain. Science 384, eadh9979 (2024).

61. Lee, B. C., Tward, D. J., Mitra, P. P. & Miller, M. I. On variational solutions for whole brain serial-section histology using a Sobolev prior in the computational anatomy random orbit model. PLOS Comput. Biol. 14, e1006610 (2018).

62. Avants, B. B., Epstein, C. L., Grossman, M. & Gee, J. C. Symmetric diffeomorphic image registration with cross-correlation: evaluating automated labeling of elderly and neurodegenerative brain. Med. Image Anal. 12, 26–41 (2008).

63. Hintiryan, H. et al. The mouse cortico-striatal projectome. Nat. Neurosci. 19, 1100–1114 (2016).

64. Bird, A. D. & Cuntz, H. Dissecting Sholl Analysis into Its Functional Components. Cell Rep. 27, 3081–3096.e5 (2019).

65. Smith, J. H. et al. How neurons exploit fractal geometry to optimize their network connectivity. Sci. Rep. 11, 2332 (2021).

66. Cortes, C. & Vapnik, V. Support-vector networks. Mach. Learn. 20, 273–297 (1995).

67. Alexander, G. E., DeLong, M. R. & Strick, P. L. Parallel Organization of Functionally Segregated Circuits Linking Basal Ganglia and Cortex. Annu. Rev. Neurosci. 9, 357–381 (1986).

68. Zingg, B. et al. Neural Networks of the Mouse Neocortex. Cell 156, 1096–1111 (2014).

69. Hintiryan, H. et al. Connectivity characterization of the mouse basolateral amygdalar complex. Nat. Commun. 12, 2859 (2021).

70. Hoover, J. E. & Strick, P. L. Multiple Output Channels in the Basal Ganglia. Science 259, 819–821 (1993).

71. Nanda, S., Das, R., Cox, D. N. & Ascoli, G. A. Structural Plasticity in Dendrites: Developmental Neurogenetics, Morphological Reconstructions, and Computational Modeling. in Neurobiological and Psychological Aspects of Brain Recovery (ed. Petrosini, L.) 1–34 (Springer International Publishing, Cham, 2017). doi:10.1007/978-3-319-52067-4_1.

72. Scheibel, M. E., Lindsay, R. D., Tomiyasu, U. & Scheibel, A. B. Progressive dendritic changes in aging human cortex. Exp. Neurol. 47, 392–403 (1975).

73. Geinisman, Y., Bondareff, W. & Dodge, J. T. Dendritic atrophy in the dentate gyrus of the senescent rat. Am. J. Anat. 152, 321–329 (1978).

74. Deutch, A. Y., Colbran, R. J. & Winder, D. J. Striatal plasticity and medium spiny neuron dendritic remodeling in parkinsonism. Parkinsonism Relat. Disord. 13 **Suppl 3**, S251–258 (2007).

75. Wang, N. et al. Msh3 and Pms1 Set Neuronal CAG-repeat Migration Rate to Drive Selective Striatal and Cortical Pathogenesis in HD Mice. Preprint at 10.1101/2024.07.09.602815 (2024).

76. Zhang, M. et al. Molecularly defined and spatially resolved cell atlas of the whole mouse brain. Nature 624, 343–354 (2023).

77. Liu, H. et al. Single-cell DNA methylome and 3D multi-omic atlas of the adult mouse brain. Nature 624, 366–377 (2023).

78. Zu, S. et al. Single-cell analysis of chromatin accessibility in the adult mouse brain. Nature 624, 378–389 (2023).

79. Zeng, H. What is a cell type and how to define it? Cell 185, 2739–2755 (2022).

80. Gouwens, N. W. et al. Classification of electrophysiological and morphological neuron types in the mouse visual cortex. Nat. Neurosci. 22, 1182–1195 (2019).

81. Migliore, M. & Shepherd, G. M. An integrated approach to classifying neuronal phenotypes. Nat. Rev. Neurosci. 6, 810–818 (2005).

82. Polavaram, S., Gillette, T. A., Parekh, R. & Ascoli, G. A. Statistical analysis and data mining of digital reconstructions of dendritic morphologies. Front. Neuroanat. 8, (2014).

83. Crittenden, J. R. & Graybiel, A. M. Basal Ganglia Disorders Associated with Imbalances in the Striatal Striosome and Matrix Compartments. Front. Neuroanat. 5, (2011).

84. Han, X. et al. Whole human-brain mapping of single cortical neurons for profiling morphological diversity and stereotypy. Sci. Adv. 9, eadf3771 (2023).

85. Peng, H., et al. Full-Spectrum Neuronal Diversity and Stereotypy through Whole Brain Morphometry. Preprint at 10.21203/rs.3.rs-3146034/v1 (2023).

86. Südhof, T. C. Synaptic Neurexin Complexes: A Molecular Code for the Logic of Neural Circuits. Cell 171, 745–769 (2017).

87. Scott, E. K. & Luo, L. How do dendrites take their shape? Nat. Neurosci. 4, 359–365 (2001).

88. Indersmitten, T., Tran, C. H., Cepeda, C. & Levine, M. S. Altered excitatory and inhibitory inputs to striatal medium-sized spiny neurons and cortical pyramidal neurons in the Q175 mouse model of Huntington’s disease. J. Neurophysiol. 113, 2953–2966 (2015).

89. Ronneberger, O., Fischer, P. & Brox, T. U-Net: Convolutional Networks for Biomedical Image Segmentation. Preprint at 10.48550/ARXIV.1505.04597 (2015).

90. Dempster, A. P., Laird, N. M. & Rubin, D. B. Maximum Likelihood from Incomplete Data Via the *EM* Algorithm. J. R. Stat. Soc. Ser. B Stat. Methodol. 39, 1–22 (1977).

91. Tward, D. et al. Diffeomorphic Registration With Intensity Transformation and Missing Data: Application to 3D Digital Pathology of Alzheimer’s Disease. Front. Neurosci. 14, 52 (2020).

92. Beg, M. F., Miller, M. I., Trouvé, A. & Younes, L. Computing Large Deformation Metric Mappings via Geodesic Flows of Diffeomorphisms. Int. J. Comput. Vis. 61, 139–157 (2005).

93. Stouffer, K. M., Witter, M. P., Tward, D. J. & Miller, M. I. Projective diffeomorphic mapping of molecular digital pathology with tissue MRI. Commun. Eng. 1, 44 (2022).

94. Tward, D. J. An Optical Flow Based Left-Invariant Metric for Natural Gradient Descent in Affine Image Registration. Front. Appl. Math. Stat. 7, 718607 (2021).

95. Breiman, L. Random Forests. Mach. Learn. 45, 5–32 (2001).

96. Abadi, M., et al. TensorFlow: A system for large-scale machine learning. Preprint at 10.48550/ARXIV.1605.08695 (2016).

97. Abdi, H. & Williams, L. J. Principal component analysis. WIREs Comput. Stat. 2, 433–459 (2010).

