## Supplementary Table 3 for "Dendritome Mapping Unveils Spatial Organization of Striatal D1/D2-Neuron Morphology"

CPx.y.z

CP = caudoputamen,

x = rostrocaudal level of CP (r, rostral; i, intermediate; c, caudal)

y = community at level x (e.g., dm, dorsomedial; dl, dorsolateral; etc.)

z = domain in community y (e.g., intermedial dorsal)

CPr.m	rostral CP, medial community

CPr.imd	rostral CP, intermediate dorsal community

CPr.imv	rostral CP, intermediate ventral community

CPr.l.ls	rostral CP, lateral community, lateral strip domain

CPr.l.vm	rostral CP, lateral community, ventromedial domain

CPi.dl.d	intermediate CP, dorsolateral community, dorsal domain (trunk/torso)

CPi.dl.imd	intermediate CP, dorsolateral community, intermedial dorsal domain (hind limb)

CPi.vl.imv	intermediate CP, ventrolateral community, intermedial ventral domain (forelimb)

CPi.vl.v	intermediate CP, ventrolateral community, ventral domain (inner mouth)

CPi.vl.vt	intermediate CP, ventrolateral community, ventral tip domain (face)

CPi.vl.cvl	intermediate CP, ventrolateral community, central ventrolateral domain

CPi.vm.v	intermediate CP, ventromedial community, ventral domain

CPi.vm.vm	intermediate CP, ventromedial community, ventromedial domain

CPi.vm.cvm	intermediate CP, ventromedial community, central ventromedial domain

CPi.dm.d	intermediate CP, dorsomedial community, dorsal domain

CPi.dm.dl	intermediate CP, dorsomedial community, dorsolateral domain

CPi.dm.dm	intermediate CP, dorsomedial community, dorsomedial domain

CPi.dm.im	intermediate CP, dorsomedial community, intermedial domain

CPi.dm.cd	intermediate CP, dorsomedial community, central dorsal

CPc.d.dm	caudal CP, dorsal community, dorsomedial domain

CPc.d.dl	caudal CP, dorsal community, dorsolateral domain

CPc.d.vm	caudal CP, dorsal community, ventromedial domain

CPc.i.d	caudal CP, intermediate community, dorsal domain

CPc.i.vm	caudal CP, intermediate community, ventromedial domain

CPc.i.vl	caudal CP, intermediate community, ventrolateral domain

CPc.v.vm	caudal CP, ventral community, ventromedial domain

CPc.v.vl	caudal CP, ventral community, ventrolateral domain

CPc.ext	caudal CP, caudal extreme (‘tail’ of the caudate)

See Hintiryan et al., 2016 for more detail.
